# Decoding protein–membrane binding interfaces from surface-fingerprint-based geometric deep learning and molecular dynamics simulations

**DOI:** 10.1101/2025.10.14.682447

**Authors:** ByungUk Park, Reid C. Van Lehn

## Abstract

Predicting protein–membrane interactions is a formidable challenge due to the subtle physicochemical features that distinguish membrane-binding regions of a protein surface, as well as the scarcity of experimentally resolved membrane-bound protein conformations. Here, we present MaSIF-PMP, a geometric deep learning model that leverages molecular surface fingerprints to predict interfacial binding sites (IBSs) of peripheral membrane proteins (PMPs). MaSIF-PMP integrates geometric and chemical surface features to produce spatially resolved IBS predictions. Compared to existing models, MaSIF-PMP achieves superior performance for IBS classification, while feature ablation studies and transfer learning analyses reveal distinct determinants governing protein–membrane versus protein–protein interactions. We further show that molecular dynamics (MD) simulations can validate model predictions, refine IBS labels, and capture composition-dependent membrane binding patterns. These results establish MaSIF-PMP as an effective framework for IBS prediction and highlight the potential of incorporating conformational dynamics from MD to improve both model accuracy and biological interpretability.

## 1. Introduction

Peripheral membrane proteins (PMPs) transiently bind to the surface of lipid membranes to mediate crucial cellular processes such as signaling, trafficking, apoptosis, lipid metabolism, and immunity.^1–3^ While PMPs exhibit a variety of molecular weights and geometries, their reversible association with the membrane is dictated by common features such as the membrane lipid environment, ions like Ca^2+^, and the polarity and geometry of the protein itself.^4^ PMPs bind to lipid membranes via noncovalent interactions with residues present at a region of the PMP surface referred to as the interfacial binding site (IBS). IBSs typically have a mixture of basic and hydrophobic amino acids^5^ that mediate non-specific electrostatic interactions and penetration into the membrane, respectively. Discovering and characterizing these IBSs is important for understanding biological mechanisms and uncovering novel sites for therapeutic intervention by modulating PMP–membrane interactions.^6–15^ However, identifying IBSs is challenging since their physicochemical properties, such as the hydropathy of IBS residues, are similar to nonbinding regions of the PMP surface.^16^ Available PMP structures are also often determined from soluble states since stabilizing membrane-bound states is difficult with contemporary experimental techniques, which impairs identification of IBSs.^7, 9, 16–18^ Alternatively, mechanistic studies of PMPs have been performed using molecular dynamics (MD) simulations, but the time required to sample PMP–membrane interactions and identify the corresponding IBS is tremendously large using all-atom MD simulations.^19, 20^ New methods are needed to rapidly characterize IBSs in order to guide the discovery of PMP-targeting drugs, the design of biosensors with enhanced binding specificity and sensitivity, and mechanistic analysis of PMP–membrane interactions.

Recently, advances in machine learning have greatly improved the accuracy and accessibility of protein structure prediction and design. Significant progress has been made in modeling protein interactions with diverse species, including other proteins, nucleic acids, small molecules, and metal ions.^21–34^ Machine learning models thus have the potential to rapidly screen large sets of PMPs to identify IBSs based on protein sequence or structural features. However, most existing models remain limited to predicting interactions between a protein and a single target ligand or other soluble species as opposed to interactions with a membrane surface. Models that have been developed to identify IBSs have primarily sought to classify individual residues as binding or non-binding.^35–38^ Such approaches lack explicit information about protein surface regions and their associated chemical properties that mediate protein–membrane interactions. One surface-centered machine learning method, referred to as molecular surface interaction fingerprinting (MaSIF)^39^, has emerged as a versatile approach to predict varied protein interactions by representing the surface of a biomolecule *(e.g*., protein, peptide, ligand) as a set of patches, each comprising numerical descriptors of geometric and chemical features. This approach has demonstrated wide applicability across diverse protein interactions (*e.g.*, with small molecules and protein) and has enabled the *de novo* design of peptides that bind to specific protein surfaces.^39, 40^ A MaSIF-based approach has also successfully designed peptides that target novel binding sites, or “neosurfaces”, that arise from protein–ligand complexes, thus expanding the target space for drug discovery.^41, 42^ These findings highlight the unique generalizability of surface fingerprint descriptors for predicting molecular interactions.

In this study, we introduce MaSIF-PMP as a geometric deep learning model that extends the MaSIF approach to predict IBSs. Because protein–membrane interactions are governed by a subtle interplay of geometric and chemical surface features, including curvature, electrostatic potentials, hydrophobicity, and hydrogen-bonding patterns^19^, that also influences protein–protein interactions, we hypothesized that MaSIF descriptors could be applied for IBS prediction. By training on a data set of nearly 1,200 PMPs, we show that MaSIF-PMP identifies IBSs with comparable accuracy to prior predictions of protein–protein interactions and outperforms state-of-the-art IBS prediction methods on a held-out test set. Unlike existing residue-based IBS prediction methods, the model generates surface-level predictions, enabling a spatially resolved landscape of potential binding regions. We further show that all-atom MD simulations can complement MaSIF-PMP predictions by validating model predictions and generating robust IBS labels. These results showcase MaSIF-PMP as a potentially powerful method for identifying IBSs, advancing understanding of protein–membrane interactions, and enabling the design of drug-like molecules to target PMPs.

## 2. Results

### 2.1. Predicting interfacial binding sites using surface fingerprints

We developed MaSIF-PMP as a geometric deep learning approach that operates on PMP molecular surface features to identify IBSs. Figure 1 presents an overview of MaSIF-PMP, including surface feature preprocessing, model architecture, and interface labeling strategy. Each protein surface is discretized into a series of points and five geometric and chemical surface features are computed for each point. Points are then assigned to a series of overlapping patches of fixed geodesic radius, with one patch centered on each point. Points within each patch are further assigned geodesic polar coordinates with respect to the patch center (Fig. 1a). The features and coordinates of the points within each patch are mapped onto learned soft polar grids, producing numerical arrays that are processed through a series of convolutional neural network (CNN) layers to yield a surface fingerprint for that patch (Fig. 1b). The resulting surface fingerprint is passed to multilayer perceptron (MLP) layers to output a score, ranging from zero to one, with higher scores indicating greater confidence that the patch is part of an IBS. Computing scores for all patches permits analysis of the likelihood of finding an IBS for all regions across the protein surface. Additional details on the surface featurization are included in the *Methods*.

**Figure 1.**
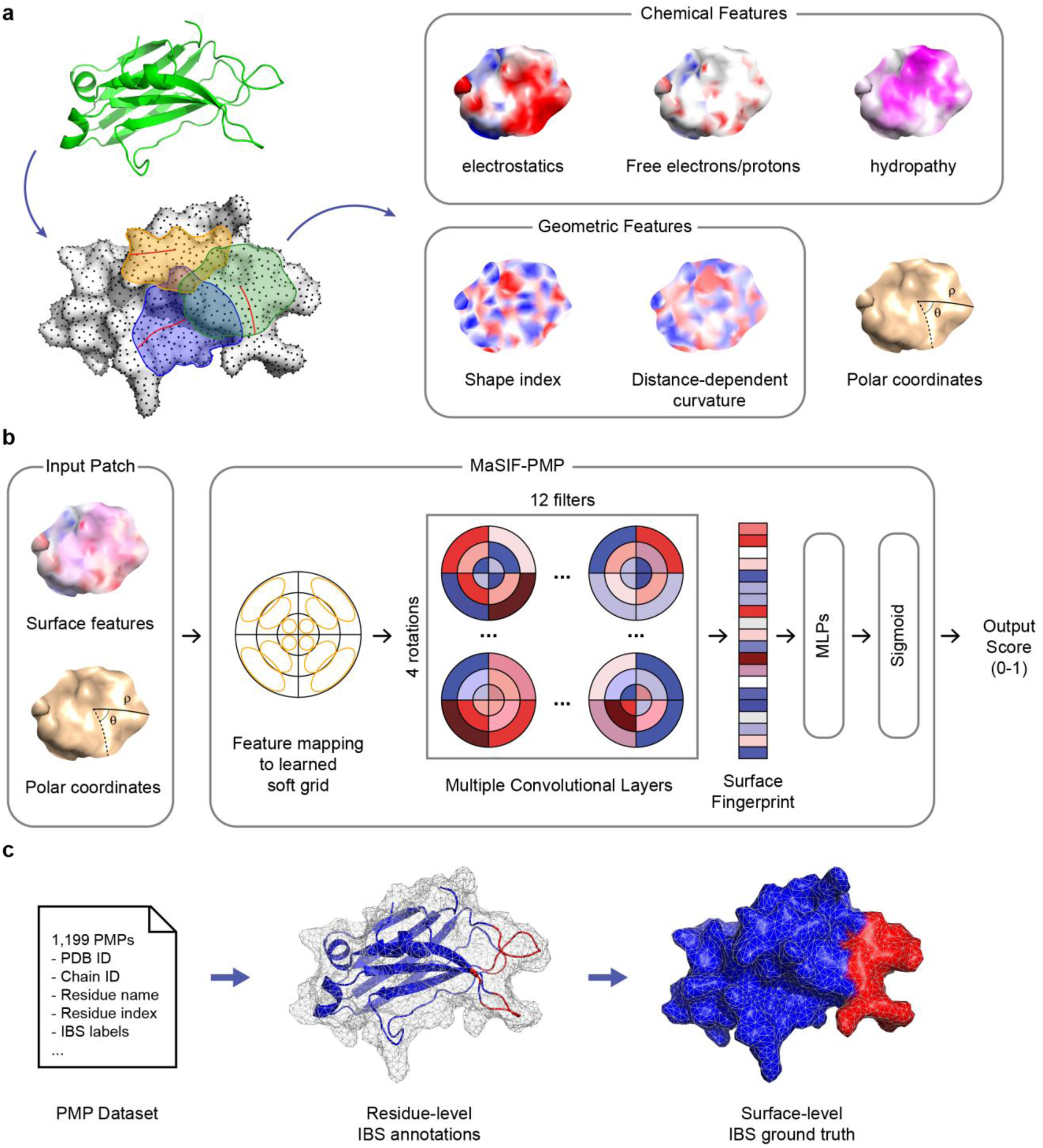
Overview of MaSIF-PMP approach. **(a)** Protein molecular surfaces are generated from atomic coordinates obtained from the Protein Data Bank and discretized into overlapping patches of fixed geodesic radius. Five geometric and chemical surface features, along with geodesic polar coordinates, are computed for each surface point.^39^ **(b)** Model architecture consisting of three convolutional layers that process each patch and output a score for the patch center. Scores range from zero to one, with higher scores indicating greater confidence that the patch is part of an interfacial binding site (IBS). **(c)** Ground-truth labels are assigned to surface points based on residue-level IBS annotations provided in a literature dataset of 1,199 PMPs.^16^ IBS-labeled residues and points are colored red, whereas non-IBS regions are colored blue.

To train MaSIF-PMP, we identified an existing dataset of 1,199 experimentally determined PMP structures with IBS annotations inferred via structural homology.^16^ Ground-truth IBS labels for model training and evaluation were derived from residue-specific annotations provided in the dataset and mapped to corresponding surface points (Fig. 1c). Specifically, each surface point was assigned the name and index of its nearest residue, allowing IBS labels to be mapped from the corresponding dataset annotations. Henceforth, we use the term “residue-level” when labeling individual amino-acid residues as being part of the IBS and the term “surface-level” when labeling individual surface points (and corresponding patches) as being part of the IBS. We split the surface-level labeled structures into a training set with 1,059 structures and test set with 130 structures that were held out during model training. Proteins were split by superfamily to preserve structural diversity, which was particularly important because the IBS labeling strategy for this data set assumes that binding patterns are conserved within a superfamily.^16^

We trained MaSIF-PMP for 50 epochs using static crystal structures obtained from the RCSB Protein Data Bank.^43^ The model at each epoch was saved based on the mean receiver operating characteristic area under the curve (ROC AUC) across a 105-protein validation set, which was set aside from the training set while maintaining similar superfamily distributions. ROC AUC is a threshold-independent metric ranging from 0 (perfect misclassification) to 0.5 (random) to 1 (perfect classification) and provides a more robust assessment of binary classifiers than metrics such as accuracy or precision (Supplementary Fig. 5). For each protein, the per-protein ROC AUC was computed by comparing surface-level predictions with ground-truth labels, and the mean plateaued at epoch 48 (Supplementary Fig. 3). On the 130-protein test set, MaSIF-PMP achieved a median per-protein ROC AUC of 0.78 for surface-level predictions (Fig. 2a), which is comparable to the performance of MaSIF when predicting protein–protein interaction sites (referred to as MaSIF-site).^39^ The most accurate MaSIF-PMP test set prediction was for the perforin C2 domain, which achieved a ROC AUC of 1.0 with predicted IBS scores closely matching ground-truth labels (Fig. 2b).

**Figure 2.**
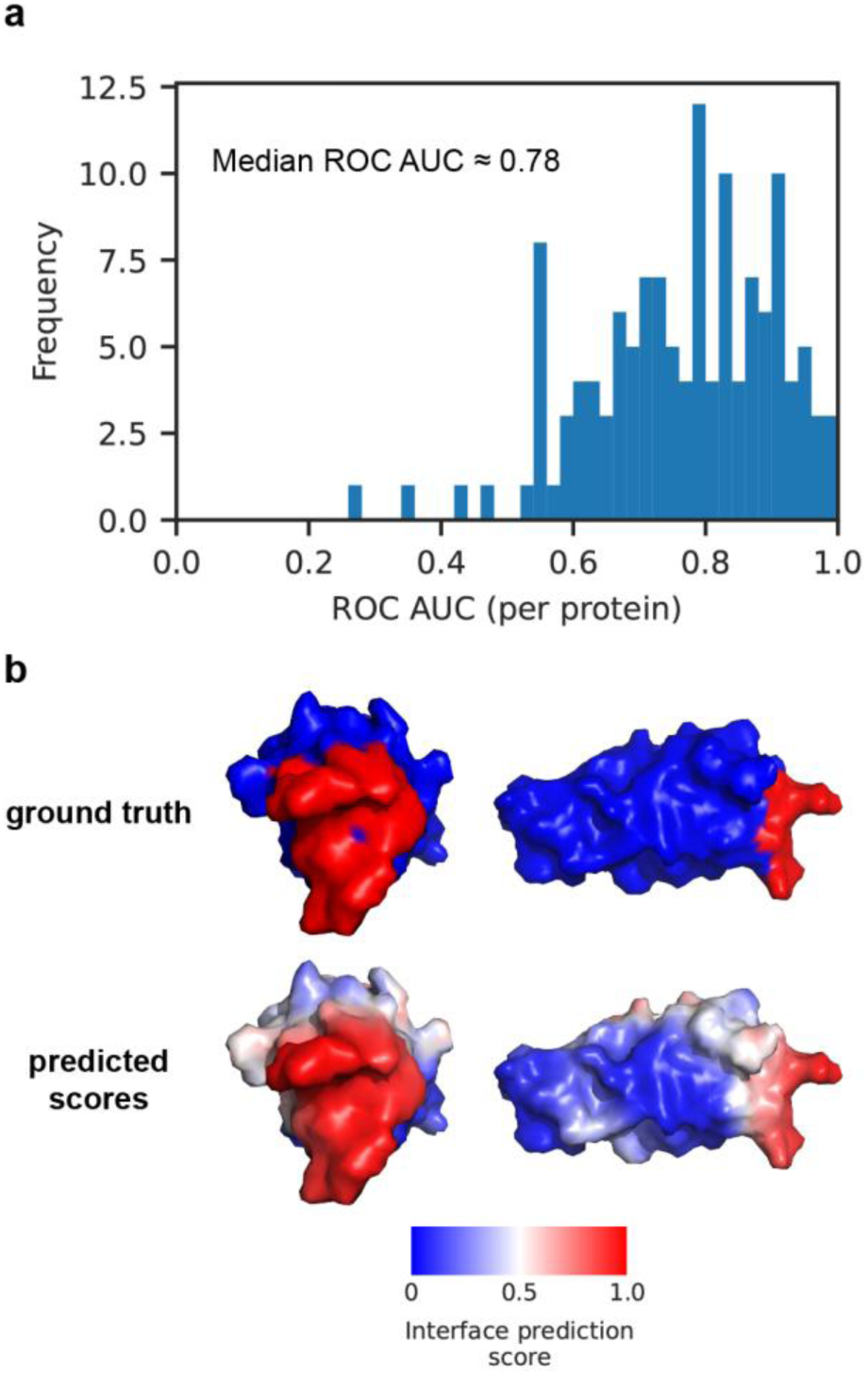
Prediction accuracy of MaSIF-PMP for a 130-protein test set. **(a)** Distribution of per-protein ROC AUC scores. Each score is computed for an entire protein by comparing surface-level predicted scores to ground-truth labels. **(b)** Snapshots of ground-truth labels and surface-level predicted scores by MaSIF-PMP for the top-performing PMP, perforin C2 domain (PDB ID: 4Y1T, chain A). IBS scores range from 0 (for nonbinding regions) to 1 (for the IBS) and are visualized using a blue-to-red color scale, with the color assigned to the point at the center of each labeled patch.

### 2.2. Benchmarking MaSIF-PMP against alternative methods

To further assess the prediction performance of MaSIF-PMP, we conducted a benchmark comparison against state-of-the-art tools for IBS prediction. Most existing methods, including DREAMM^36^, PPM3^37^, and MODA^38^, produce residue-level predictions, while PMIpred^35^, a physics-informed model, generates predictions at the level of 15-residue segments. In contrast, MaSIF-PMP outputs surface-level predictions. To compare between models, we mapped MaSIF-PMP predictions from surface- to residue-level and likewise mapped predictions from other models from residue- to surface-level (Supplementary Fig. 5 and Note). This benchmark was performed using 21 PMPs from the PMIpred^35^ benchmark set with corresponding subunit structures and their ground-truth IBS residue labels. For consistency, we updated the labels of three PMPs (PDB IDs: 1JSS, 2RSG, 1LN1) from the benchmark set—which were also included in our test set but annotated with broader IBS regions—to use the same labels as our test set. The same MaSIF-PMP model introduced in the previous section was used for the benchmark without retraining. Performance was evaluated using ROC AUC and the Matthews correlation coefficient (MCC; see Equation S5), calculated over surface points and surface-exposed residues only. The MCC was included as an evaluation metric due to its suitability for imbalanced classification tasks; in our case, non-binding regions greatly outnumber IBS regions. MCC values range from -1 to +1, with +1 indicating perfect classification, 0 corresponding to random guessing, and -1 indicating perfectly incorrect classification.

MaSIF-PMP outperformed all other predictors in both per-protein residue-level ROC AUC and MCC. Kernel density estimates of the ROC AUC (Fig. 3a) and MCC (Fig. 3b) show a clear shift to higher values for MaSIF-PMP compared to other predictors, reflecting improved overall performance. Consistently, MaSIF-PMP achieved higher median per-protein, residue-level ROC AUC and MCC values (Fig. 3c) as well as per-protein surface-level evaluations (Supplementary Fig. 6-7). Together, these results demonstrate that MaSIF-PMP surpasses existing IBS predictors across multiple metrics, underscoring its accuracy and robustness in handling imbalanced labels.

**Figure 3.**
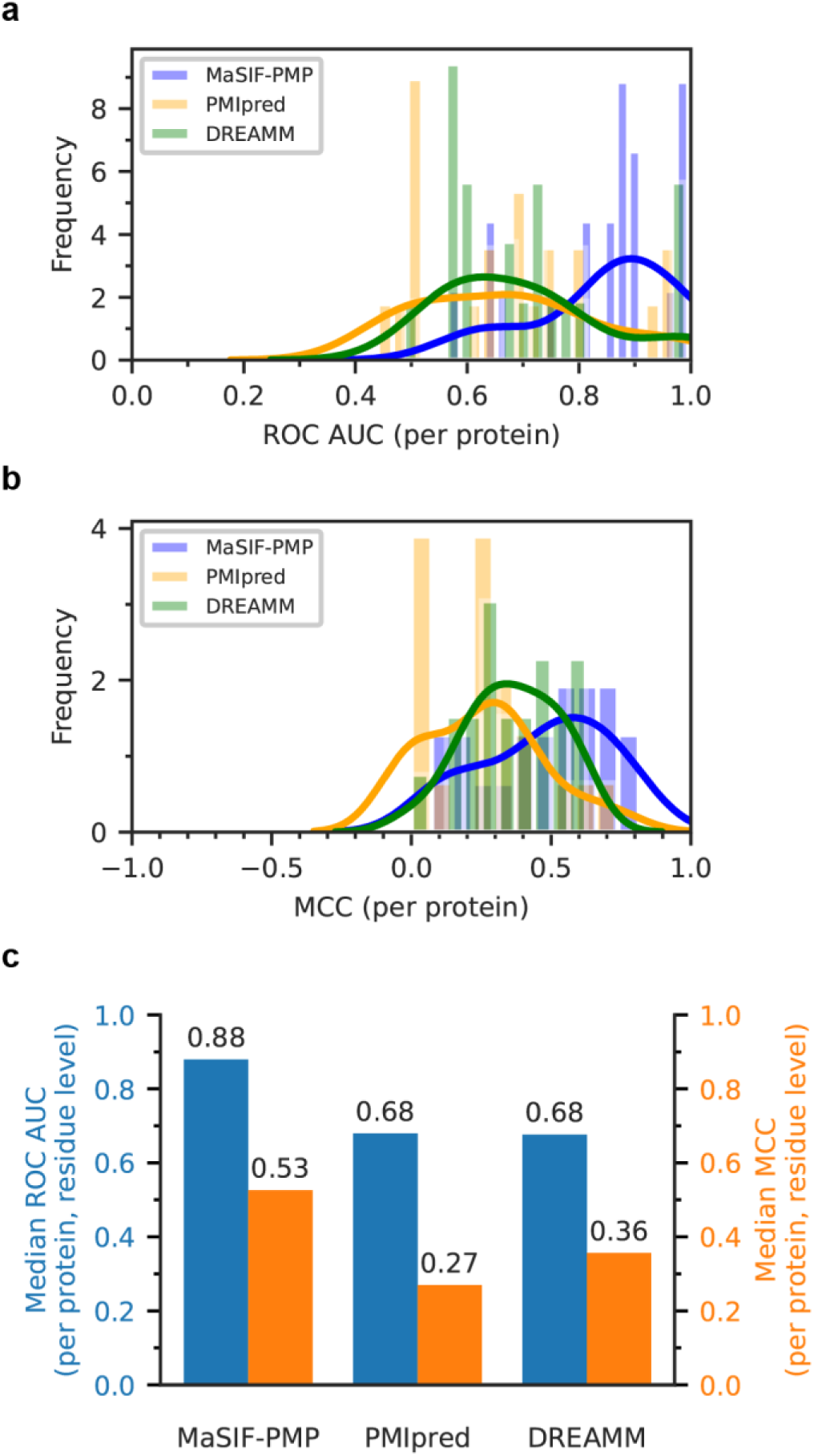
Benchmark comparison of MaSIF-PMP against other IBS predictors assessed by the ROC AUC and MCC for 21 single-chain PMPs. Distributions of per-protein, residue-level **(a)** ROC AUC and **(b)** MCC predicted by MaSIF-PMP, PMIpred^35^, and DREAMM^36^. Solid lines represent Gaussian kernel density estimates fit to the discrete score distributions. **(c)** Comparison of MaSIF-PMP with other IBS predictors on the benchmark proteins. Results are reported as the median ROC AUC and median MCC per protein, evaluated on a per-residue basis to ensure comparability across predictors.

Surface-level comparisons (Fig. 4) further demonstrate that MaSIF-PMP predicts broader interaction interfaces than PMIpred and DREAMM. In contrast to the binary outputs of other predictors, MaSIF-PMP generates continuous prediction scores that can be visualized as surface color gradients. Given that protein–membrane interactions arise from extended protein–membrane surface contacts, our surface-level approach provides a more realistic representation of binding interfaces than residue-level predictions that are limited to a small subset of anchoring residues. For instance, in the case of sphingomyelinase C (Fig. 4c), the residue-level predictors achieved higher ROC AUC and MCC values because the annotated interface constitutes only a small portion of the protein surface. However, the broader IBS predicted by MaSIF-PMP likely reflects regions that approach the membrane during interaction and, thus, contribute to protein–membrane interactions, as further discussed in the context of MD simulations below. MaSIF-PMP can also produce binary outputs by applying a defined threshold, enabling deployment in the same manner as other predictors to designate specific surface patches as IBS regions (Supplementary Fig. 9). Together, these results show that MaSIF-PMP not only outperforms existing IBS predictors but also provides a more holistic view of the protein–membrane interface and the underlying interaction process.

**Figure 4.**
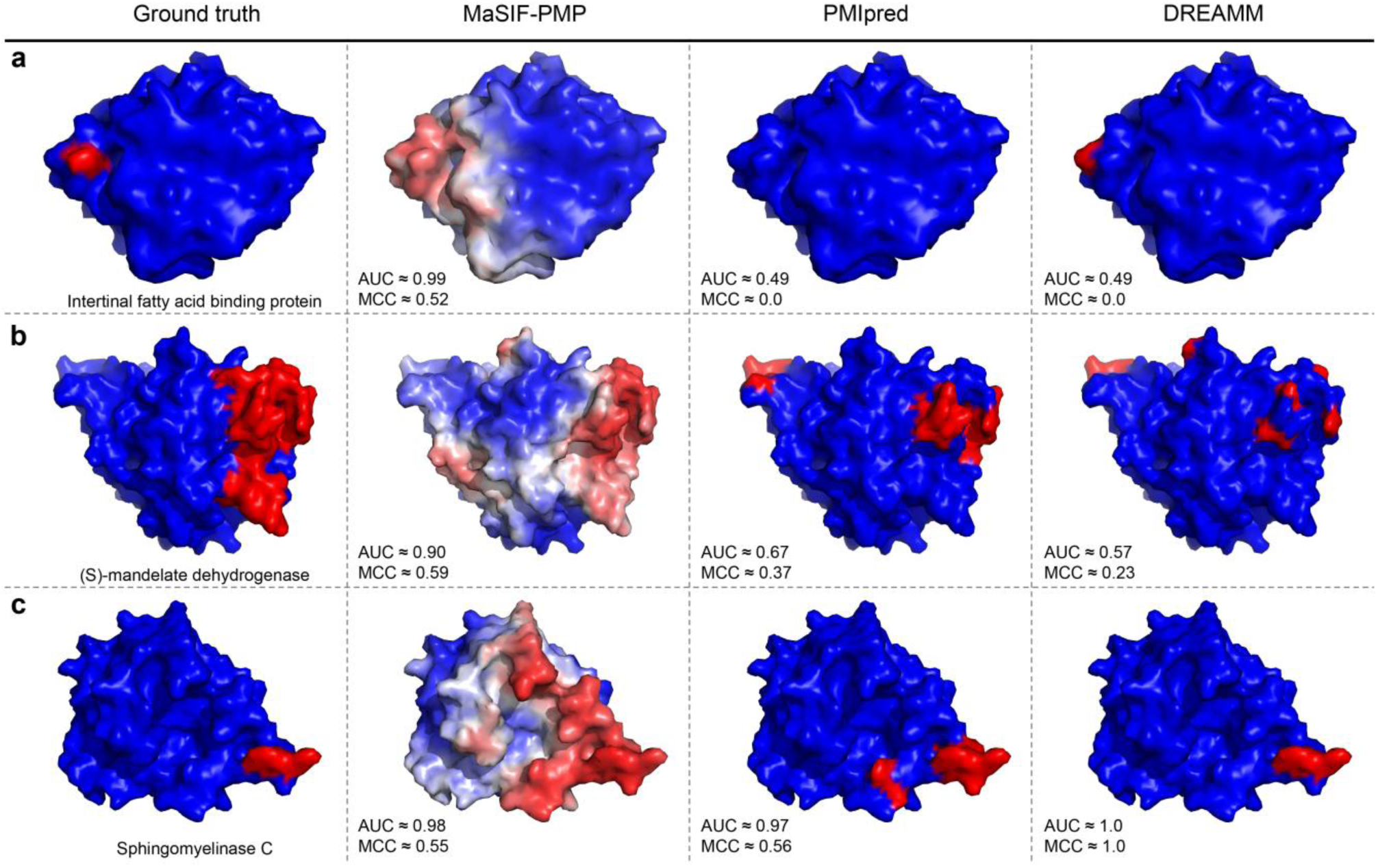
Visualization of ground-truth and predicted IBSs for three PMPs selected from the benchmark test set. Each column shows PMP molecular surfaces with colors indicating binary labels (0 for non-binding surface points or 1 for the IBS) for the ground-truth, PMIpred, and DREAMM columns and continuous interface prediction scores (from 0 to 1) for MaSIF-PMP following the color scheme in Fig. 2. ROC AUC and MCC values are computed based on surface-level predictions for all three models. **(a)** Intestinal fatty acid binding protein (PDB ID: 3AKM). **(b)** (S)-mandelate dehydrogenase (PDB ID: 6BFG). **(c)** Sphingomyelinase C (PDB ID: 2DDR).

### 2.3. Analysis of interplay between surface features in protein–membrane interactions

To assess the relative contributions of surface features to IBS predictions, we performed an ablation study by conducting five-fold cross-validation on the complete 1,199 PMP dataset using models trained with different subsets of surface features. We hypothesized that the network trained on the most critical features would yield the highest median ROC AUC scores. As shown in Fig. 5a, among models trained on individual feature types, the network using only geometric features (shape index and distance-dependent curvature) achieved the highest ROC AUC of 0.72, followed by models trained using only hydropathy, electrostatic potential, and hydrogen bonding features. The prediction performance of the network trained with two geometric features was comparable to that of the model trained with all three chemical features, underscoring the dominant role of geometric information in defining IBS patterns. This finding contrasts with protein–protein interaction predictions obtained using MaSIF-site, for which single chemical features outperformed geometric features.^39^ Furthermore, we observed a cumulative improvement in predictive accuracy as additional features were added as input for training, with the highest performance achieved when all five surface features were included (Supplementary Fig. 10). These results indicate that geometric features are more critical than individual chemical features for IBS prediction in PMPs, and that integrating all five features maximizes the prediction performance, reflecting the subtle interplay of geometric and chemical determinants in protein–membrane interactions.

**Figure 5.**
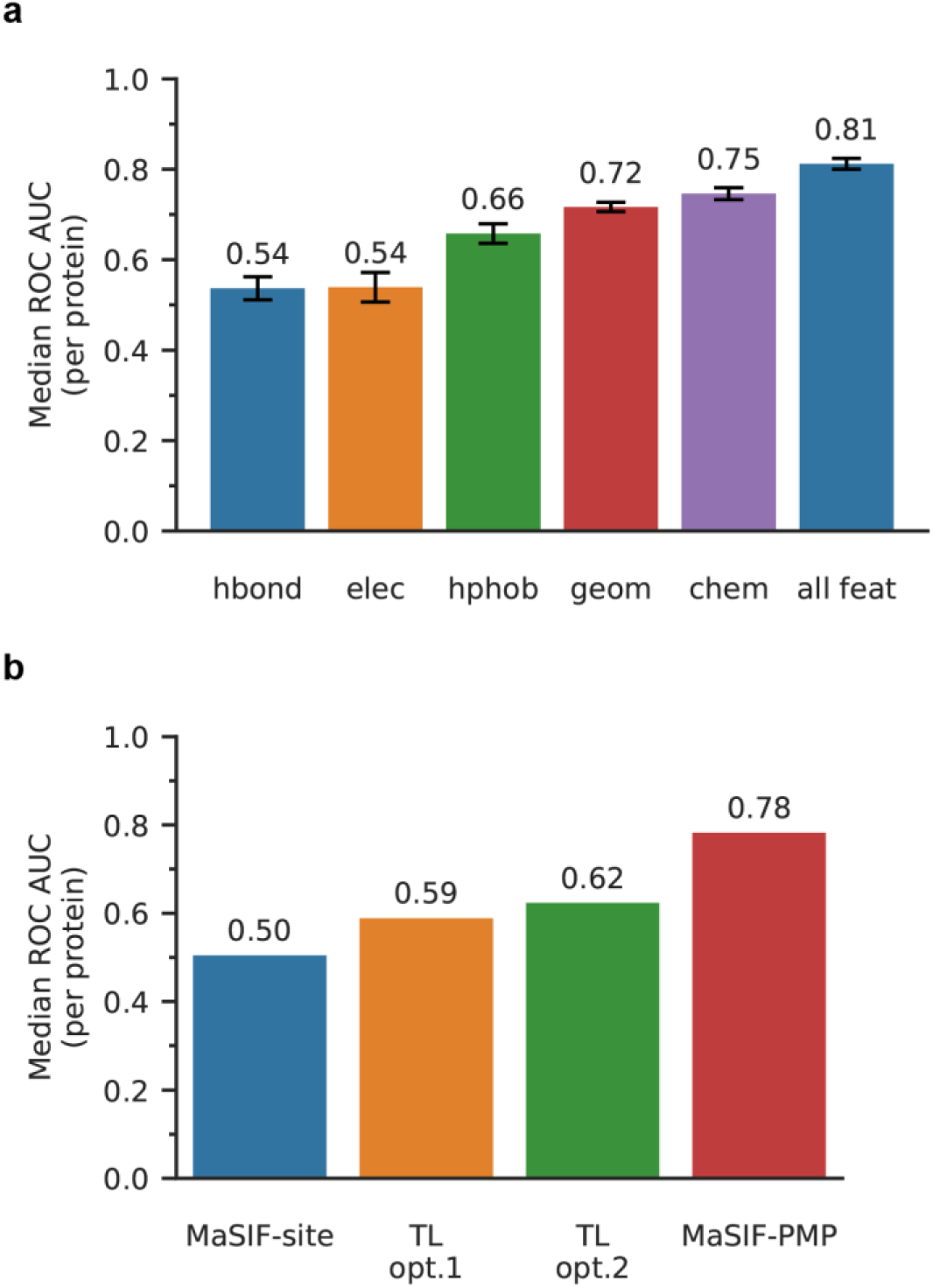
Comparisons of different approaches for MaSIF-PMP training. **(a)** 5-fold cross-validation with models trained on different subsets of surface features: only the location of free electrons/proton donors (hbond), only electrostatic potential (elec), only hydropathy index (hphob), two geometric features (geom), all three chemical features (chem), and all five chemical and geometric features (all feat) **(b)** Transfer learning of the MaSIF-site model^39^, which is trained on protein–protein interactions, to IBS predictions. Each column indicates the median ROC AUC for the 130-PMP test set with MaSIF-site, MaSIF-PMP with transfer learning strategy 1 (TL opt.1) and strategy 2 (TL opt.2), and MaSIF-PMP without transfer learning.

Because surface fingerprint descriptors have demonstrated material-agnostic applications in predicting diverse protein–ligand interactions^39–42^, we next tested whether we could derive models from the MaSIF-site model, which is trained for protein–protein interactions, to improve IBS predictions via transfer learning. That is, we sought to determine whether the latent space of surface fingerprints for protein–protein interactions is sufficiently similar to the latent space of fingerprints for IBS predictions so that the extensive protein–protein interaction data used to train MaSIF-site could be leveraged to improve MaSIF-PMP. Two transfer learning (TL) strategies were employed: (i) freezing convolutional layers and retraining only a deeper fully connected network (TL opt.1) using the PMP training data, or (ii) freezing convolutional layers and adding new convolutional layers for retraining (TL opt.2). Fig. 5b compares model accuracy on the 130 PMPs test set for MaSIF-site (*i.e.*, a model trained on protein–protein interactions without any retraining), TL opt.1, TL opt.2, and MaSIF-PMP (*i.e.*, a model trained on labeled IBS without any transfer learning). MaSIF-site yielded a median ROC AUC of 0.50, indicating an essentially random guess when applied for IBS prediction. Both TL strategies produced only modest improvements and did not approach the prediction accuracy of MaSIF-PMP. These results, together with the ablation study presented in Fig. 5a, emphasize the fundamental differences in how surface features govern protein–protein versus protein–membrane interactions.

### 2.4. Data augmentation using MD simulations of PMPs in solution

MaSIF-PMP demonstrated strong performance in predicting IBSs using surface features from static crystal structures. We next asked whether incorporating conformational dynamics from MD simulations could further improve performance because previous studies have shown that PMPs often undergo conformational changes upon membrane binding.^20^ Crystal structures typically represent soluble-state conformations, which may differ substantially from the membrane-bound state and thus fail to capture IBS-relevant structural features. MD simulations provide an effective means to explore conformational dynamics and potentially identify conformations more relevant to membrane interactions^20, 44–49^, but their computational cost increases with the number of simulated particles, and the required bilayer dimensions and complexity can vary widely between PMPs.

To circumvent these challenges, we hypothesized that unbiased MD simulations of proteins in aqueous solution may capture conformational dynamics relevant to their membrane-bound states, thereby enabling efficient sampling of important conformations at a lower computational cost than simulating protein–membrane interactions. To test this hypothesis, we performed high-throughput MD^50^ simulations of PMPs from the training set in aqueous solution and sampled representative conformations from trajectories using CLoNe.^51^ We then augmented the training dataset with these MD-derived conformations to evaluate their impact on prediction for the test set. The model trained on MD-augmented data had mean and median per-protein ROC AUC scores of 0.77 and 0.78, respectively, compared to 0.76 and 0.78 for the model trained only on crystal structure (Supplementary Fig. 12). This nearly identical performance indicates that conformations sampled from aqueous-phase simulations provide limited improvement to IBS predictions, likely because they are not representative of membrane-bound states.

### 2.5. Case studies using MD simulations with the HMMM model

Although MD simulations of PMPs in solution were not effective for improving MaSIF-PMP predictions, we next sought to evaluate whether MD simulations could (i) validate predicted IBSs for PMPs with experimental annotations, (ii) generate IBS labels for PMPs lacking experimental annotations, and (iii) enhance prediction accuracy by sampling PMP conformations relevant to the membrane-bound state. We thus evaluated case studies using MD simulations with the highly mobile membrane mimetic (HMMM) model. The HMMM model accelerates lateral lipid diffusion by truncating full-length lipids and replacing the bilayer hydrophobic core with an organic solvent region.^52^ This approach enables sampling of protein– membrane interactions at reduced computational cost while preserving atomic-level interaction details.

We first performed MD simulations of α-tocopherol transfer protein (α-TTP), a well-characterized PMP not included in our training dataset (Fig. 6). α-TTP is a cytosolic liver protein that undergoes conformational changes upon membrane binding, notably involving distortion of its N-terminal domain toward the membrane plane^53, 54^, but previous MD studies^47^ did not explicitly identify membrane-contacting residues at its IBS. We prepared systems with the protein positioned ∼1 nm above the HMMM membrane and performed unbiased simulations to sample protein–membrane interactions. IBS regions were defined from the membrane-bound conformations at the end of the trajectories using a 0.5 nm distance threshold between heavy atoms of protein side chains and lipid atoms at the membrane surface. We further performed replica simulations with multiple orientations of α-TTP relative to the HMMM membrane to assess the reliability of the MD-based IBS labels. Although a small number of organic solvent molecules escaped from the HMMM bilayer (Fig. 6a), this effect reflects the natural partitioning of the solvent between lipid and solution^55^ and does not affect IBS predictions. The residues in contact with the membrane were consistent across the trajectory in each replica simulation, leading to robust HMMM-derived IBS labels (Supplementary Fig. 13). Using the α-TTP crystal structure as input and HMMM-derived labels from the final membrane-bound simulation configuration as ground-truth, MaSIF-PMP achieved a ROC AUC of 0.81, which remained comparable (0.75) when the HMMM-derived membrane-bound conformation was used instead (Fig. 6b). α-TTP exhibited stable membrane interactions in specific orientations, with high overlap among IBS labels derived from different replicas, and MaSIF-PMP’s prediction accuracy on the crystal structure was comparable whether IBS labels were taken from individual replicas or from a consensus IBS identified based on time-averaging over multiple simulation configurations (Supplementary Fig. 13). These results highlight the robustness and reliability of the HMMM-based IBS labels when compared to MaSIF-PMP predictions.

**Figure 6.**
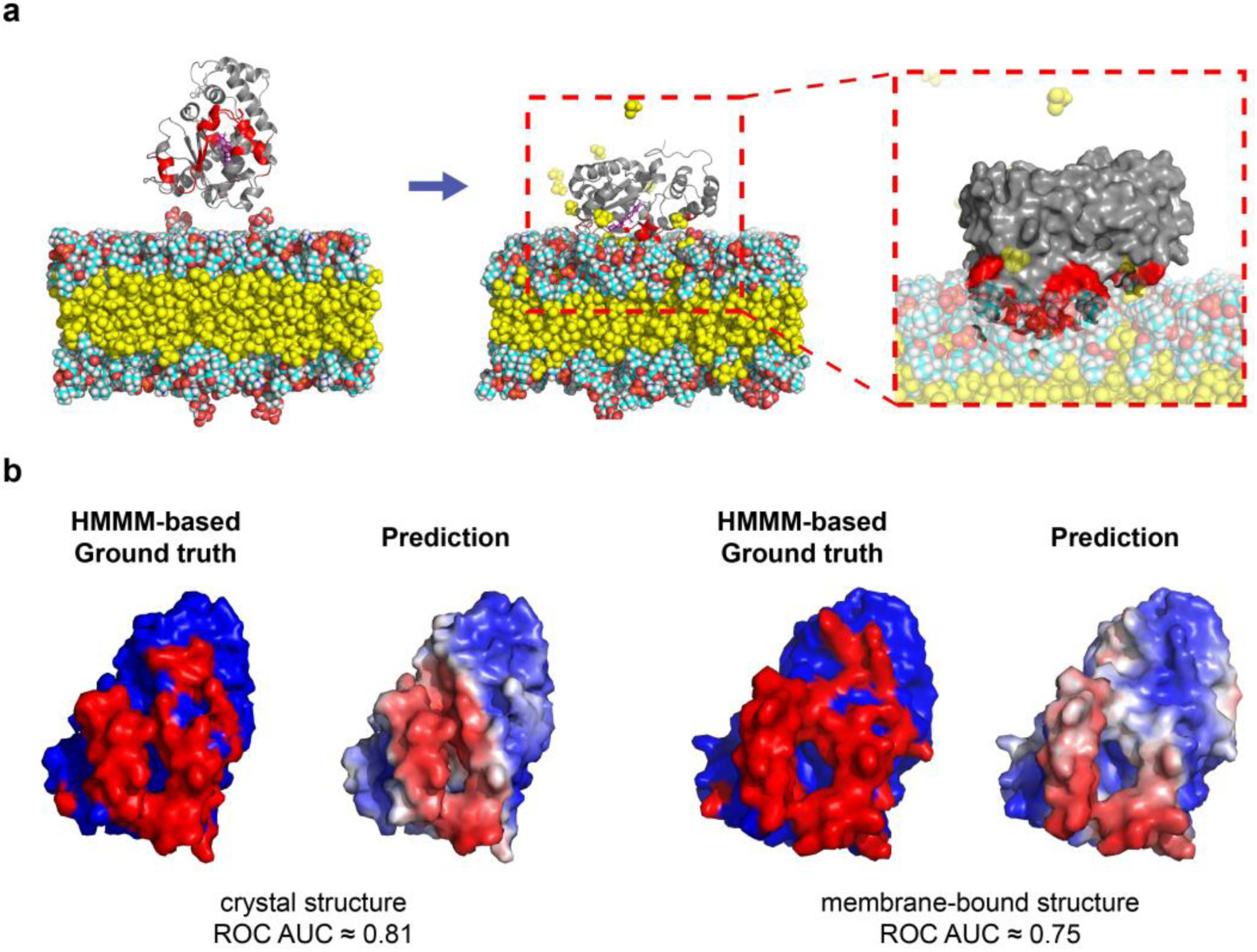
HMMM MD simulations of PMP-membrane binding and comparison to MaSIF-PMP predictions. **(a)** Representative snapshots of α-TTP after 100 ns HMMM MD simulation. The IBS contacting the membrane is highlighted in red, with an enlarged snapshot of the protein in contact with the membrane within the red dashed box. The HMMM membrane is shown in a van der Waals representation with carbon atoms in cyan, hydrogen atoms in white, oxygen atoms in red, nitrogen atoms in blue, phosphorus atoms in orange, and organic solvent 1,1-dichloroethane (DCLE) in yellow. α-tocopherol, a bound ligand in complex with α-TTP, is in purple. α-TTP is drawn in a ribbon representation in gray, with the surface shown in the enlarged image. Water molecules and ions are not shown to aid visualization. **(b)** Comparison of ground-truth IBS labels determined by HMMM MD and MaSIF-PMP predictions for α-TTP using either the crystal structure or the membrane-bound conformation obtained from HMMM MD.

We observed similar patterns in a second case study with the oxysterol-binding protein homologue (Osh4), a PMP that undergoes structural rearrangement in which six initially separated domains reorganize into a composite IBS with the anionic trans-Golgi network (TGN) membrane.^20, 45, 56^ We performed HMMM simulations using lipids that mimic a TGN membrane and sampled a membrane-bound conformation within 100 ns; prior MD simulations required ∼300 ns to observe membrane binding using full-length lipid simulations^20^, demonstrating the computational efficiency of the HMMM approach. HMMM-predicted IBS labels overlapped with the labels from the previous studies^20, 45^, although they encompassed fewer residues within the IBS regions. Using IBS labels from the literature studies^20, 45^, MaSIF-PMP yielded ROC AUC values of 0.63 for the crystal structure and 0.64 for the membrane-bound conformation (Supplementary Fig. 14). In contrast, with HMMM-predicted IBS labels, MaSIF-PMP attained ROC AUC values of 0.74 for the crystal structure and 0.68 for the membrane-bound conformation (Supplementary Fig. 14), underscoring the robustness and accuracy of HMMM-derived IBS labels compared to both prior annotations and MaSIF-PMP predictions.

Next, we performed HMMM MD simulations of PMPs with IBSs that were poorly predicted in the MaSIF-PMP test set to investigate whether poor prediction performance stemmed from limitations in the dataset’s IBS annotations—namely, due to homology-based labels that lacked direct experimental confirmation.^16^ We selected two such proteins: phospholipase A2 (PDB ID: 1OZY) and glycosyl hydrolase (PDB ID: 4LPL), which yielded ROC AUC scores of 0.54 and 0.42, respectively, when their crystal structures were used as model input. HMMM simulations revealed membrane composition-dependent binding modes for phospholipase A2, with distinct IBS regions emerging in anionic versus zwitterionic membranes (Fig. 7a, Supplementary Fig. 15). The anionic membrane yielded more consistent consensus IBSs across replica simulations with different initial PMP orientations, suggesting a more favorable and stable binding mode (Supplementary Fig. 15). IBS labels combined from these replicas (referred to as the “union IBS”) captured the dynamic nature of the protein–membrane interface (Supplementary Fig. 15 and Note). Notably, MaSIF-PMP predictions have a higher ROC AUC when compared to the anionic HMMM-derived union IBS than with either the original dataset labels or the zwitterionic HMMM-derived union IBS (Fig. 7b), reflecting the model’s ability to identify preferred, membrane-composition-specific binding interfaces. A similar trend was observed for glycosyl hydrolase, although membrane composition had less influence on binding mode (Supplementary Fig. 16, Fig. 7c). These findings suggest that the omission of membrane composition relevant to PMP binding may contribute to low prediction performance, potentially due to mislabeling in the original dataset when using homology-based labeling, and highlight the potential of HMMM simulations to refine poorly predicted labels at a manageable computational cost.

**Figure 7.**
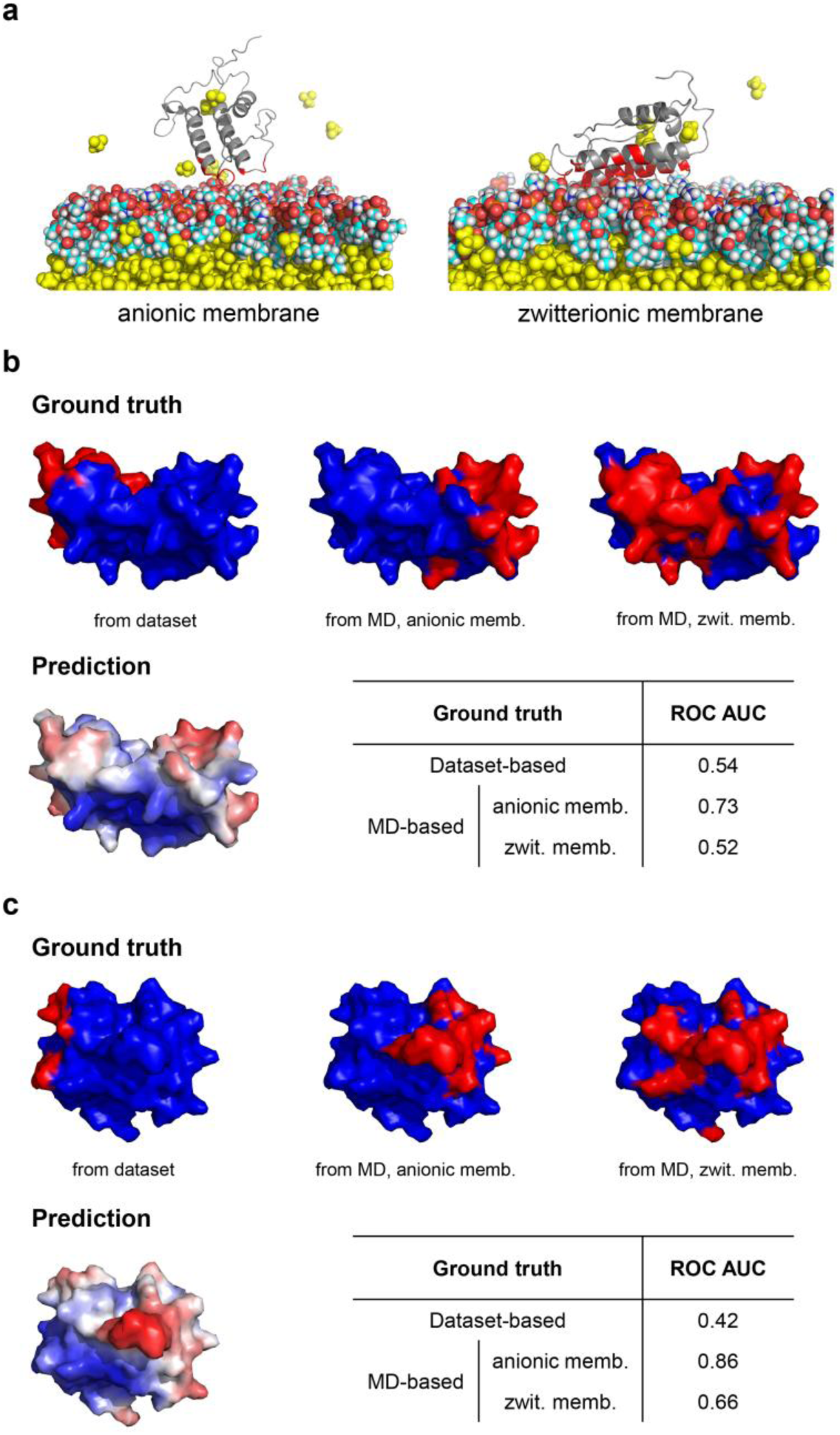
HMMM MD as a tool to generate interface labels for proteins lacking experimental annotations. **(a)** Snapshots of phospholipase A2 (PDB ID: 1OZY) in contact with different types of membranes sampled from HMMM MD simulations. IBSs for each membrane are highlighted in red. Color schemes follow the same definition used in Figure 6a. **(b-c)** Snapshots of ground-truth IBS labels (based on dataset annotations and HMMM MD simulations) and predicted IBS scores for **(b)** phospholipase A2 (PDB ID: 1OZY) and **(c)** glycosyl hydrolase (PDB ID: 4LPL). Tables show corresponding ROC AUC values computed with different ground-truth label types. “anionic memb.” refers to an anionic membrane, while “zwit. memb.” refers to zwitterionic membrane.

Together, these case studies demonstrate that MD simulations with the HMMM model are suitable for capturing conformational dynamics in membrane binding and refining IBS annotations. While membrane-bound conformations from HMMM yielded reasonable but slightly lower prediction accuracy than crystal structures, these results suggest that MaSIF-PMP predictions are consistent with HMMM labels when using either crystal structures or MD-derived conformations as input, and that improved selection of representative membrane-bound states could enhance model agreement.

## 3. Discussion

In this work, we developed MaSIF-PMP, a geometric deep learning model that leverages surface fingerprints to predict IBSs on PMPs. By encoding both geometric and chemical surface features into numerical descriptors, the model learns to discriminate between IBS and non-IBS patches following training on a large PMP dataset. Our results demonstrate that surface fingerprints serve as robust, material-agnostic descriptors, extending their applicability beyond protein–protein interactions to protein–membrane systems, while revealing distinct latent feature spaces underlying each interaction type. In addition, MD simulations with the HMMM model proved valuable for validating predictions, capturing distinct membrane-bound conformations of proteins depending on membrane composition, and refining IBS labels at manageable computational cost.

Despite these advances, several limitations of our approach should be acknowledged. The current convolutional neural network architecture, which incorporates rotational data augmentation and processes single proteins with at least 1,000 patches per batch, is memory intensive. As computational resources allow, training with larger surface patches, extended geodesic radii, and soft grids with more trainable Gaussian kernels may enhance prediction performance. Dataset-related limitations also remain: the model was trained on single-chain protein structures and thus does not account for cases where PMPs interact with membranes as multimers (*e.g.*, BAR proteins that bind membranes as dimers to sense curvature^57^). Furthermore, many IBS annotations of the PMP dataset were indirectly inferred based on homology rather than directly validated; generating more robust labels via MD simulations and incorporating information about specific membrane compositions could provide richer training data and enhance generalizability.

Several directions may further advance this framework in future work. Incorporating conformational flexibility into surface fingerprints, particularly geometric features, or into IBS annotations as soft labels through dynamic ensembles derived from HMMM simulations could improve predictions for PMPs undergoing structural rearrangements. Integrating such dynamic conformations, along with membrane surface features determined by lipid composition, may ultimately improve the predictive accuracy and biological relevance of surface-based models and enable the development of a framework for identifying proteins that bind to membranes with distinct compositions that is analogous to methods to identify protein-protein interaction partners.^39^ Replacing certain descriptors with more efficient alternatives, such as topological data analysis features, could accelerate preprocessing and enable training with more extensive surface information. Finally, predicted IBSs of PMPs implicated in diseases through dysregulated membrane interactions may serve as therapeutic targets by enabling explicit modeling of PMP–ligand interactions.

## 4. Methods Dataset

PMPs were obtained from a previously published dataset comprising 1,199 experimentally determined PMP structures, each annotated with constituent amino acids, IBS labels, and structural features.^16^ Ten PMPs from the dataset were excluded from the training and testing sets; some were omitted due to extensive unmodeled residues in the central region of their sequences, which resulted in misleading surface representations, while others lacked IBS annotations entirely (Supplementary Table 1). The training and testing sets consisted of 1,059 and 130 proteins, respectively, following the same split ratio used in the original study of MaSIF-site.^39^ Proteins were split based on their superfamilies to maintain similar superfamily distributions between sets (Supplementary Table 2) an important consideration since the IBS labeling procedure for the dataset assumes that structurally related proteins within the same superfamily share similar IBSs. IBS annotations were primarily inferred through structural superposition, aligning PMP structures with homologous domains and transferring IBSs from annotated members supported by experimental or simulation-based evidence.^16^ This approach provided a scalable means of annotation but represents a compromise necessitated by the high computational cost of validating IBS labels via molecular dynamics simulations for all proteins in the dataset. Further details on the structural split are described in Supplementary Note.

### Model: MaSIF-PMP

MaSIF-PMP is similar to the previously developed protein interaction site prediction model, MaSIF-site^39^. Compared to MaSIF-site, our model architecture is implemented in PyTorch v2.1.2^58^ instead of Tensorflow v1^59^, PMP datasets are used for training and testing, and the method for defining interface labels for PMPs differs from the original model.

### Computation of discretized molecular surfaces

The MSMS program^60^ was used to compute all molecular surfaces in this study (density = 3.0, water radius = 1.5 Å). As MSMS generates molecular surfaces with highly irregular meshes, the resulting meshes were further regularized using PyMESH (v.0.3.1)^61^ at a resolution of 1.0 Å. Consequently, each protein’s molecular surface is represented as a discretized triangulated mesh (Supplementary Fig. 1).

### Decomposition of protein surfaces into overlapping radial patches

For each point on the discretized protein surface mesh, we extracted a radial patch with a geodesic radius of 9 Å to capture local surface information.^62^ Each patch was limited to a maximum of 100 mesh points. This choice of a 9 Å radius with a 100-point cap was made to reduce memory requirements, thereby allowing the use of multiple convolutional layers, which are critical for accurate interface prediction.^39^ The geodesic distances—defined as distances along the protein surface—required to define these patches were computed using the Dijkstra algorithm^63^.

### Computation of geodesic polar coordinates

After extracting surface patches from a protein, our geometric deep-learning pipeline computes radial and angular coordinates for the mesh points within each patch. Representing local surface features in this geodesic polar coordinate system allows the model to map the patch onto a 2D plane. The radial coordinate is determined using the Dijkstra algorithm, which computes the geodesic distance from the patch center to each point. To determine the angular coordinate, pairwise geodesic distances between points within the patch are first computed, and multidimensional scaling (MDS)^64^, as implemented in scikit-learn^65^, is then used to project the points onto a 2D plane. A random direction in this plane is selected as the 0° reference axis, and the angle of each point relative to this axis is computed following previously described procedures^39^.

### Geometric and chemical surface features

Each point within a patch of the computed molecular surface was assigned an array of two geometric features (shape index^66^ and distance-dependent curvature^62^) and three chemical features (hydropathy index^67^, Poisson-Boltzmann electrostatic potential^68^, and hydrogen bond potential^69, 70^). These features are identical to those used in MaSIF-site.^39^ Details on each feature and how to compute are described in Supplementary Note.

### Definition of interfacial binding site points in a protein surface

Using the residue-level IBS annotations of PMPs in the dataset, we defined the ground-truth labels for the protein surface mesh points. Since each mesh point carries name and index attributes of its nearest residue, IBS labels could be assigned directly based on the corresponding dataset annotation for the nearest residue.

### Geometric deep learning on a learned soft polar grid

Geometric deep learning extends deep learning techniques, such as convolutional neural networks (CNNs)^71^, to non-Euclidean data structures like protein surfaces. In this work, we employed a soft polar grid—a system of Gaussian kernels defined in a local geodesic polar coordinate system that act as soft, overlapping pixels—as described previously for MaSIF-site.^39^ This soft grid serves as an analogue to the sliding window used in conventional CNNs for image analysis^71^, where a window moves across the image, extracts a patch of pixels, multiplies them by corresponding learnable filter weights, and sums the results. The parameters of the Gaussian kernels are themselves learnable, enabling the network to adaptively capture local surface patterns.^39, 72^ Once the mapping is complete, a traditional CNN layer is applied together with angular max pooling, ensuring that the resulting fingerprint vectors are invariant to the randomly assigned polar coordinates of geodesic patches. Further methodological details are provided in Supplementary Note, Supplementary Fig. 2 and Refs. 39, 72.

### Neural network, cost function and training optimization

The neural network architecture consists of three convolutional layers with learnable soft grids (Supplementary Fig. 2). The optimal number of convolutional layers was determined through a systematic evaluation by incrementally increasing the number of layers (Supplementary Fig. 3). The network takes as input the full protein surface decomposed into overlapping patches with a radius of 9.0 Å. The patch radius was chosen to balance surface coverage with memory efficiency, thereby enabling the use of deeper convolutional layers. Each patch is mapped onto learned grids with three radial bins and four angular bins. The network outputs an IBS score between 0 and 1 for the center point of each patch. During training, each protein was treated as a single batch, and the network was optimized using an Adam optimizer^73^ with a sigmoid cross-entropy loss function. Because non-IBS points greatly outnumber IBS points in most proteins, negative samples were randomly subsampled to balance the dataset and ensure an equal number of positive and negative samples. Training was performed on an NVIDIA L40 GPU for 50 epochs (approximately 1.55 hours), with all proteins in the training set processed once per epoch. Model performance was evaluated using the per protein ROC AUC metric. 10% of the training data was set aside as a validation set to monitor model performance. The model was saved whenever the validation ROC AUC improved, with the final saved model occurring at epoch 48. Beyond epoch 50, the validation ROC AUC reached a plateau, indicating that 48 epochs were sufficient for the network to converge (Supplementary Fig. 3a).

### 5-fold cross-validation with feature masking

We performed 5-fold cross-validation using the PMP dataset to evaluate the contribution of individual surface features to IBS characterization. For each fold, models were systematically trained with specific features masked, such that the masked features were excluded from training. All experiments were conducted for 50 training epochs.

### Transfer Learning

We performed two different strategies for transfer learning of MaSIF-site, which is trained on protein–protein interactions, to MaSIF-PMP. First, we froze the updated parameters of convolutional layers of MaSIF-site and replaced the final fully connected networks with deeper ones (FC128, FC64, FC4, FC2). Second, we froze the same convolutional layers then added new 3 convolutional layers so that only the parameters of additional layers and final fully connected network block are updated during training on PMP dataset. Schematics and further details of each structure are provided in Supplementary Note and Supplementary Fig. 11. Training on protein–protein interactions was performed under the same conditions as MaSIF-site^39^ whereas training on the PMP dataset followed the conditions described for MaSIF-PMP in the *Methods*.

### Highly Mobile Membrane-Mimetic (HMMM) Simulations

#### HMMM model

The highly mobile membrane-mimetic (HMMM) model accelerates the lateral diffusion of lipid molecules by one to two orders of magnitude while preserving atomic-level interaction details by replacing the hydrophobic core with an organic solvent region and truncating full-length lipids.^52^ This approach is advantageous for all-atom protein–membrane interactions simulations, as conventional full-length lipid bilayers exhibit slow membrane reorganization dynamics. HMMM systems were prepared using the CHARMM-GUI HMMM Builder.^55^ Default parameters were used for the terminal acyl carbon number (6), proteins were protonated assuming pH 7.0, standard charged termini were applied (N-terminus: -NH ^+^; C-terminus: -COO^-^), 1,1-dichloroethane (DCLE) was used as an organic solvent, and systems were neutralized with NaCl counterions in explicit water. All simulations employed the CHARMM36m force field and TIP3P water model.^74–76^ Systems were energy-minimized and equilibrated under NPT conditions using the CHARMM-GUI–generated simulation protocols. Production simulations of 100 ns were performed at constant NPT, maintaining a temperature of 310.15 K with a velocity-rescale thermostat and a pressure of 1 bar using a semi-isotropic stochastic cell-rescaling barostat. Further details are provided in Supplementary Note.

#### System preparation for case studies

For α-tocopherol transfer protein (α-TTP), we adopted conditions from a previous MD study^47^, using the same crystal structure (PDB ID: 3W67), lipid composition, and simulations under NPT conditions at 300 K. For additional case studies involving PMPs with low prediction performance, HMMM simulations were performed with two representative membrane types (anionic and zwitterionic) under NPT conditions at 310.15 K and 1 bar. In all systems, proteins were initially placed ∼1 nm above the HMMM membrane surface using different orientations for different replicas. For oxysterol-binding homologue (Osh4), we used the experimentally determined structure (PDB ID: 1ZHZ)^56^, which has also been employed in previous computational studies of Osh4–membrane interactions using all-atom simulations.^20, 45^ HMMM membranes with anionic lipid composition were prepared to match the conditions of these prior studies. Simulations were performed under NPT conditions at 303.15 K, consistent with the experimental and computational setups reported previously.^20, 45^ Further details of system setup and parameters are provided in Supplementary Note.

#### IBS labels determination from HMMM simulations

The last simulation configuration of each trajectory for which the protein and HMMM membrane were in contact was used to define IBS labels. Protein residues with heavy atoms within 0.5 nm of the heavy atoms of truncated lipids on the membrane surface were designated as IBS residues. Details on defining consensus and union IBS labels using trajectories of replica simulations are described in a Supplementary Note.

## Supplementary Information Description

The supplementary information includes additional information on MaSIF-PMP parameters and surface feature calculations, additional validation tests and comparisons, and details on simulation methods and parameters. The Supplementary Information includes 16 Supplementary Figures and 5 Supplementary Tables.

## Acknowledgements

This material is based upon work supported by the National Science Foundation under Grant No. CBET-2346683. This work used the Center for High Throughput Computing (CHTC, doi:10.21231/GNT1-HW21).

