## Supplemental Information for "Decoding protein–membrane binding interfaces from surface-fingerprint-based geometric deep learning and molecular dynamics simulations"

### Supplementary Information

ByungUk Park[1] and Reid C. Van Lehn[1,2,*]

[1]Department of Chemical and Biological Engineering and [2]Department of Chemistry

University of Wisconsin−Madison, Madison, Wisconsin 53706, United States

### PDBs excluded from training and test sets

The PMP dataset[1] published in 2022 originally comprised 1,199 proteins with residue-level annotations of membrane-binding and non-binding interfaces. Of these, 10 proteins were excluded from both the training and test sets, resulting in a final dataset of 1,189 proteins. Seven of these ten proteins were removed due to extensive unmodeled regions in the central part of their sequences, which led to poorly defined molecular surface representations. The remaining three proteins were excluded because they lacked true interface annotations entirely. The corresponding PDB and chain identifiers are listed in Supplementary Table 1.

**Supplementary Table 1.** PDB and chain IDs of proteins excluded from the PMP dataset used for model training and evaluation. Each entry is listed as a PDB ID followed by its chain ID, separated by an underscore.

| | **PDB_CHAIN IDs** |
|---|---|
| Unmodeled residues in the central region of the sequence | 2DYB_B, 3QBV_D, 4UXJ_H, 5CCG_E, 5KJ7_K, 3M7F_B, 3TVV_B |
| No interface annotations | 1P8J_B, 2ID4_A, 4OMD_B |

### Train–test split rationale

The curated dataset of 1,189 proteins was split into 1,059 training and 130 test proteins, following the same training-to-test ratio used in the original MaSIF-site study.[2] To preserve structural diversity and ensure generalizability, the split was performed such that both sets maintained similar distributions of proteins across different superfamilies. This consideration was particularly important given that the dataset's interfacial binding site (IBS) labeling strategy relies on the assumption that structurally related proteins within the same superfamily share similar membrane-binding sites. The number of proteins assigned to each superfamily in the training and test sets is detailed in Supplementary Table 2.

**Supplementary Table 2.** Distribution of proteins across superfamilies in the training and test sets. Superfamily labels correspond to membrane-targeting domains (Annexin, C1, C2, Discoidin C2, PH, PX), enzymes (PLA, PLC/D), and lipid-transfer proteins (START).

| | Training set | Test set | Total |
|---|---|---|---|
| Annexin | 42 | 5 | 47 |
| C1 | 33 | 4 | 37 |
| C2 | 90 | 11 | 101 |
| Discoidin C2 | 256 | 32 | 288 |
| PH | 234 | 29 | 263 |
| PX | 41 | 5 | 46 |
| PLA | 126 | 15 | 141 |
| PLC/D | 60 | 7 | 67 |
| START | 177 | 22 | 199 |
| Total | 1059 | 130 | 1189 |

### Data preprocessing

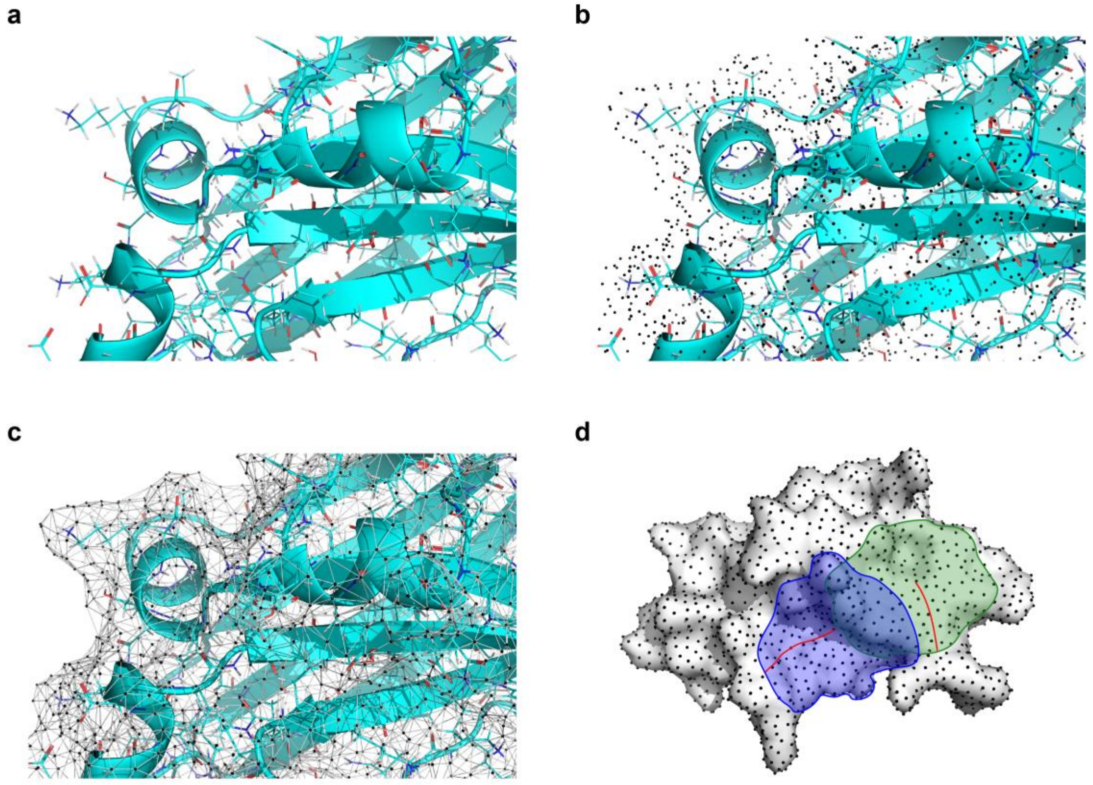


**Supplementary Figure 1.** Illustration of the decomposition of protein surfaces into overlapping radial patches. The input protein structure **(a)** is first preprocessed to generate dot surface **(b)** using the MSMS program.[3] A graph representation **(c)** is then constructed, where vertices correspond to the surface dots and edges connect neighboring vertices. This mesh is further regularized to produce a discretized triangulated surface. Each surface vertex serves as the center of radial patch **(d)**, and neighboring vertices within a fixed geodesic radius (9 Å in MaSIF-PMP) are included in the patch. Surface features of the vertices are computed and embedded into a numerical array representing the patch-level input.

### Surface feature calculations

#### Shape index

The shape index (Equation S1) describes the local geometry around each point on the surface based on curvature.[4] Its values range from −1 (highly concave) to +1 (highly convex) and are defined in terms of the principal curvatures $\kappa_1$ and $\kappa_2$, where $\kappa_1 \geq \kappa_2$ (Equation S2), as follows:

$$\text{shape index} = \frac{2}{\pi}\tan^{-1}\frac{\kappa_1 + \kappa_2}{\kappa_1 - \kappa_2} \tag{S1}$$

$$\kappa_1 = H + \sqrt{H^2 - K}, \kappa_2 = H - \sqrt{H^2 - K}, \tag{S2}$$

*H* is the vertex mean curvature and *K* is the vertex Gausssian curvature, which were both computed using PyMESH.[5]

#### Distance-dependent curvature

For each vertex within an extracted patch, the distance-dependent curvature computes a value in the range [−0.7, 0.7] to capture the relationship between the surface point's distance to the patch center and the orientation of its surface normal relative to the center point. Further details on this feature are provided in Ref. 4. While the shape index, a principal curvature-based descriptor, characterizes the local geometry of each vertex across the entire protein surface, the distance-dependent curvature quantifies curvature at the patch level using the patch center as a reference. This feature has been shown to capture complementary information to the shape index.[2, 4]

#### Poisson-Boltzmann electrostatic potential

We used PDB2PQR[6] to prepare protein structures for electrostatic calculations, and Poisson–Boltzmann electrostatics were computed using APBS[7] (v.1.5). The electrostatic potential at each vertex of the triangulated molecular surface was assigned using Multivalue, a utility within the APBS suite.[7] Charge values exceeding +30 or falling below −30 were capped at those respective limits, after which all values were normalized to a range between −1 and 1.

#### Hydrogen bond potential

The locations of free electrons and potential hydrogen bond donors on the molecular surface were computed using a hydrogen bond potential model as described in Ref. 8. Surface vertices whose nearest atom was a polar hydrogen, nitrogen, or oxygen were identified as potential hydrogen bond donors or acceptors. A value drawn from a Gaussian-shaped potential function was then assigned to each vertex based on the orientation between the relevant heavy atoms. These values range from −1 (optimal position for a hydrogen bond acceptor) to +1 (optimal position for a hydrogen bond donor).

#### Hydropathy index

Each surface point was assigned a hydropathy value based on the Kyte and Doolittle scale[9], according to the amino acid identity of the closest atom. Original values, which range from −4.5 (most hydrophilic) to +4.5 (most hydrophobic), were normalized to a range between −1 and 1.

### Geometric deep learning on protein surface using learned soft polar grid

Using learned soft polar grid on a molecular surface enables a generalization of the convolutional networks (CNN) paradigm to non-Euclidean manifolds.[10, 11] The learned soft polar grid used in this work contains $\theta$ angular bins and $\rho$ radial bins for a total of $J = \rho\theta$ bins. For each vertex in the discretized molecular surface $x$, with neighbors $N(x)$ and each vertex $y \in N(x)$, we define the coordinates $u(x, y)$ as the radial and angular coordinates of $y$ with respect to $x$. The mapping of each grid cell $j$ for feature vector $f$ and the patch centered at $x$, $D_j(x)f$, is defined as:

$$D_j(x)f = \sum_{y \in N(x)} w_j\big(u(x,y)\big)f(y)\,, j = 1, \dots, J \tag{S3}$$

where $w_j$ is a Gaussian weight function and $f(y)$ are the features at vertex $y$. In our model, 4 angular bins and 3 radial bins are used, resulting in a total of $J = 12$ Gaussian bins. To ensure rotational invariance in the neural network, we performed 4 rotations of the input patch and conducted a max-pool operation on the output.[12] Further details of using learned soft polar grid on molecular surface are described in Refs. 2, 10, 11.

### Model architecture

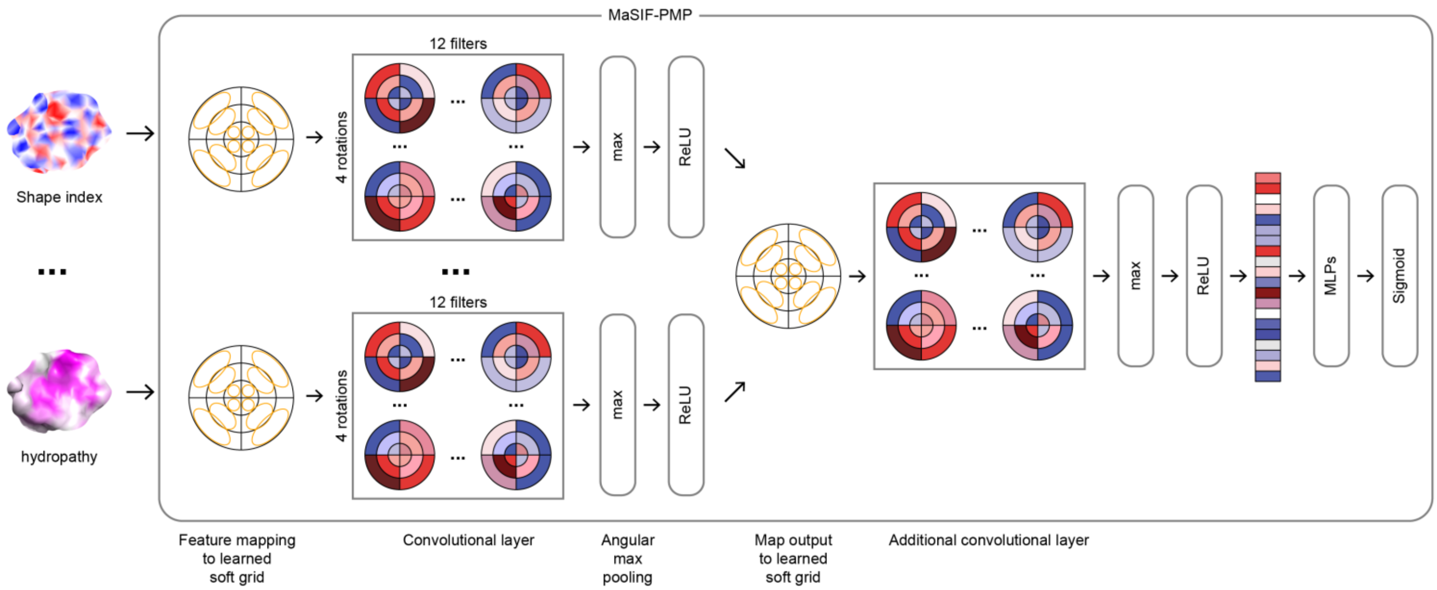


**Supplementary Figure 2.** Network architecture of MaSIF-PMP. Surface patches are processed through a series of convolutional layers, followed by multilayer perceptron (MLP) blocks, and trained using a sigmoid cross-entropy loss function.

### Evaluation of model structure for MaSIF-PMP

We evaluated the effect of incorporating layer normalization into the MaSIF-PMP architecture to improve depth stability (Supplementary Fig. 3). Layer normalization standardizes feature values per surface point, stabilizing feature scales across the diverse patches and regions on the protein. Batch normalization or instance-wise spatial normalization across points may not be adequate as each point is semantically independent and the surface point number (*i.e.*, batch size) varies depending on proteins. While layer normalization increased model stability, its inclusion led to slightly lower prediction performance compared to models without it. In particular, performance decreased when layer normalization was applied to architectures with more than five convolutional layers. For models without layer normalization, instability emerged when the number of convolutional layers exceeded five. The highest prediction performance was achieved with a five-layer architecture without layer normalization.

We also tested whether having more convolutional layers might lead to overfitting of the model by comparing two networks with different number of convolutional layers (Supplementary Fig. 4). Based on the distribution of predicted IBS scores of all surface points from test set, we observed that the network with five convolutional layers was better at classifying true negatives (non-interface points) but shows flat distributions of scores for true positives (interface points). However, the network with three convolutional layers outputted more predictions with high interface scores, indicating that it is better for distinguishing IBS from non-IBS regions despite the lower prediction accuracy compared to the model with five convolutional layers. We thus chose the network with three convolution layers, yielding the same structure as previous work on predicting protein-protein interactions (PPI) using MaSIF-site.[2]

Given that (i) depth-induced instability was not a critical issue at this configuration, (ii) the original MaSIF-site model for PPI prediction[2] was also implemented with three convolutional layers and without layer normalization, and (iii) the network with three convolutional layers yielded comparable prediction accuracy to the one with five layers (best-performing one) while better at predicting true binding interfaces, we selected the three-layer, no–layer-normalization architecture as the final model used for comparisons in the main text. Nonetheless, layer normalization may become necessary if the number of surface features used as input descriptors is substantially increased in future work.

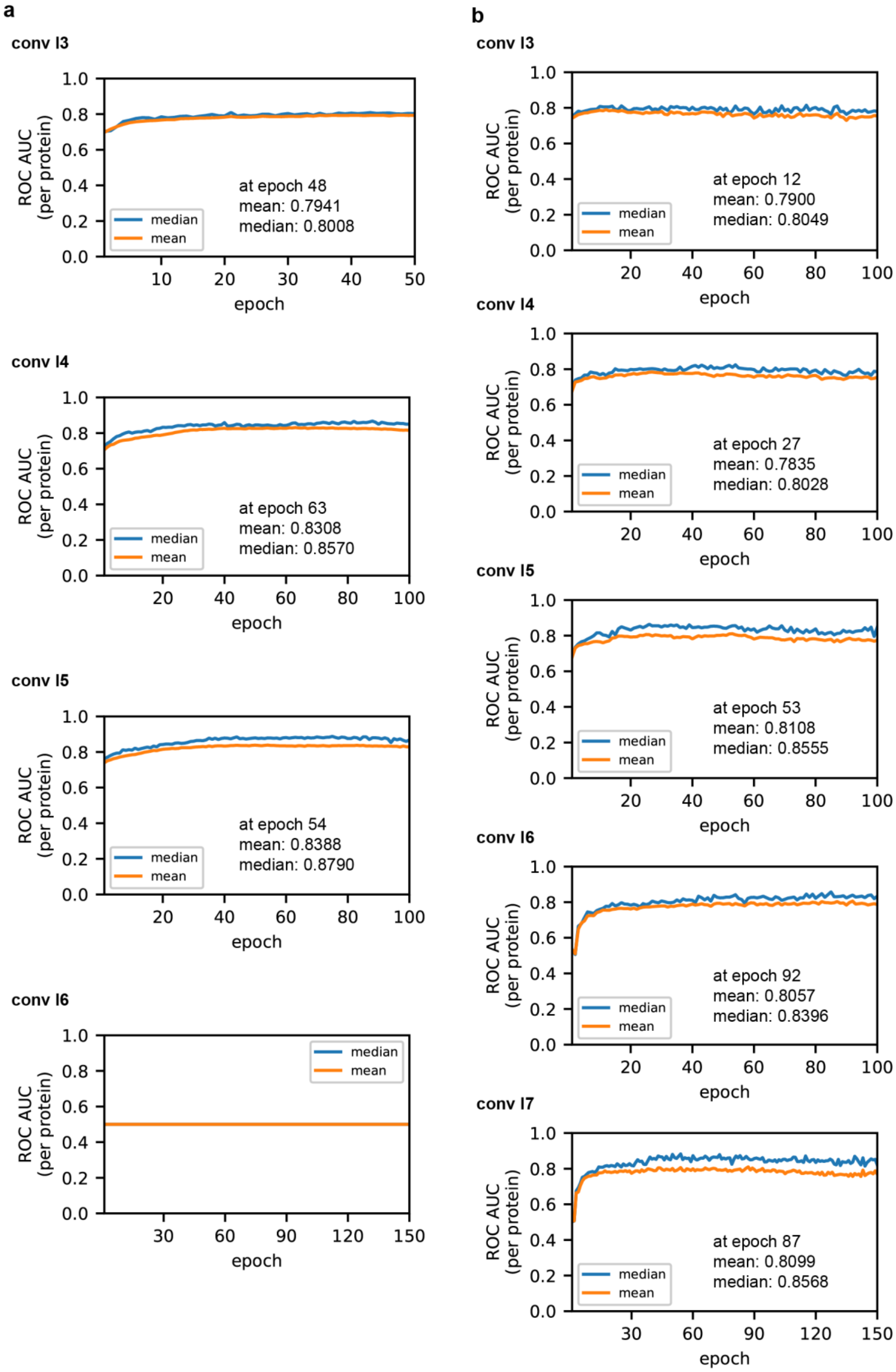
a
conv l3
at epoch 48
mean: 0.7941
median: 0.8008
conv l4
at epoch 63
mean: 0.8308
median: 0.8570
conv l5
at epoch 54
mean: 0.8388
median: 0.8790
conv l6
b
conv l3
at epoch 12
mean: 0.7900
median: 0.8049
conv l4
at epoch 27
mean: 0.7835
median: 0.8028
conv l5
at epoch 53
mean: 0.8108
median: 0.8555
conv l6
at epoch 92
mean: 0.8057
median: 0.8396
conv l7
at epoch 87
mean: 0.8099
median: 0.8568
ROC AUC (per protein)
epoch
median
mean

**Supplementary Figure 3.** Systematic evaluation of the number of convolutional layers (noted as "conv l#") of MaSIF-PMP. Mean and median of per protein ROC AUC scores for validation set were plotted as functions of epoch number. **(a)** Evaluations using models without layer normalization. The best model performance was observed when it contained five convolutional layers, while more layers introduced a depth-induced instability that led to a flat ROC AUC of 0.5 for all epochs. All tested models reached plateau in mean of per protein ROC AUC values within the tested epoch number. The inset text indicates the epoch number at which the best performing model was saved and the corresponding mean and median per-protein ROC AUC values. **(b)** Evaluations using models with layer normalization. Incorporating layer normalization addressed the depth-induced instability and model prediction performance decreased as the model used more than five convolutional layers.

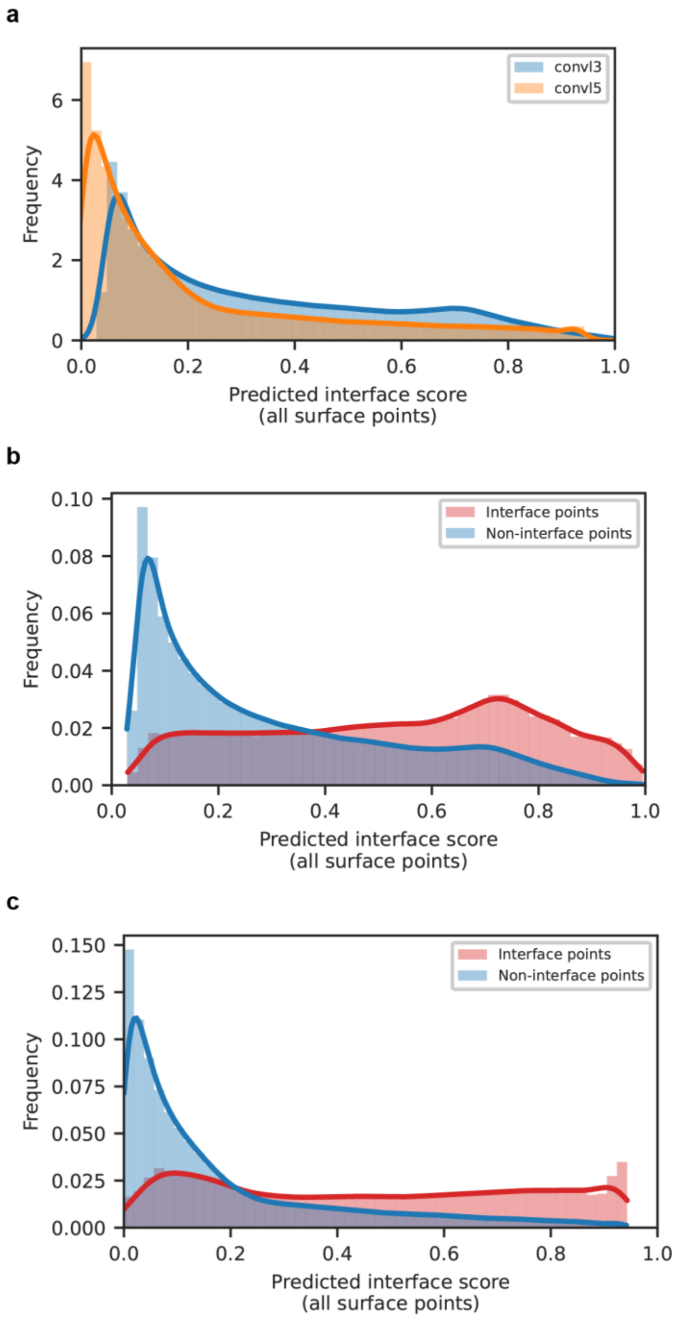


**Supplementary Figure 4.** Distribution of predicted interface scores for the test set using models with different number of convolutional layers. **(a)** Distribution of predicted scores for all surface points in the test set for different models. "convl3" indicates the model with three convolutional layers while "convl5" indicates the model with five layers. **(b-c)** Distribution of predicted interface scores of test set for true positives (red) versus true negatives (blue) for a network trained with all features. Prediction results from model trained with **(b)** three convolutional layers and **(c)** five convolutional layers.

### Comparison with other models

For the benchmark comparison, we selected two state-of-the-art IBS predictors, PMIpred[13] and DREAMM.[14] The benchmark test set was selected from the PMIpred study and includes 21 proteins with experimentally resolved structures downloaded from the RCSB PDB—19 of which are from the DREAMM test set, and 2 are known lipid packing defect sensors. To ensure consistency and comparability, AlphaFold-predicted structures were excluded. Model predictions were generated using the same PDB and chain IDs as those in the PMIpred benchmark. Ground-truth labels were also taken directly from the PMIpred benchmark, except for three proteins (PDB IDs: 1JSS, 2RSG, 1LN1). For these, we used the broader IBS annotations from our dataset, which already encompassed the original PMIpred labels but included additional interface residues.

#### Mapping surface-level predictions to residue-level

To allow direct comparison with PMIpred and DREAMM, which provide residue-level interface predictions, we mapped MaSIF-PMP's surface-level outputs to the residue level (Supplementary Fig. 5a). Since MaSIF-PMP generates predictions only at mesh points on the molecular surface, we restricted evaluation to surface-exposed residues.

We followed the same strategy used in the benchmark assessment of MaSIF-site[2], a previous model for PPI prediction, to convert surface-level predictions into residue-level scores. In particular, we used the SPPIDER definition[15] of interfacial residues, which has been extensively validated both qualitatively and quantitatively. According to this definition, interface residues are those whose solvent-excluded surface area changes by at least 5 Å$^2$ upon binding and contribute at least 4% to the total solvent-excluded interface area. We note that all calculations were performed using solvent-excluded, not solvent-accessible, surface areas for consistency with MaSIF's surface representation.

Residue-level IBS scores from MaSIF-PMP were computed by assigning each residue the maximum IBS score among all its associated surface points. Conversely, for PMIpred and DREAMM, whose outputs are at the residue-level, we mapped their predictions onto the molecular surface using the same method described in the main text (*Methods* section) for defining IBS labels on surface points.

**Metrics for comparison**

To evaluate and compare predictor performance, we used both area under curve (AUC) of the receiver operating characteristic (ROC) curve and the Matthews correlation coefficient (MCC) as evaluation metrics. ROC AUC is a threshold-independent metric and offers a more robust assessment than basic metrics such as accuracy or precision (Supplementary Fig. 5b). Specifically, given the model-predicted scores and corresponding ground-truth labels, threshold values were systematically varied from 0 to 1 to convert continuous scores into binary labels: residues with scores equal to or above the threshold were assigned positive labels and the rest were assigned negative labels. For a given threshold, a point was classified as a true positive (TP) or true negative (TN) if its predicted label matched the corresponding positive or negative ground-truth label, and as a false positive (FP) or false negative (FN) if it did not. True and false positive rates (Equation S4) were then computed for all thresholds to generate the ROC curve, and the area under this curve was used as the ROC AUC metric. ROC AUC scores range from 0 to 1, where 1 indicates a perfect model and 0.5 indicates random guessing. Higher AUC scores (*e.g.*, > 0.7) signify better model performance, demonstrating a greater ability to distinguish between positive and negative classes across all possible thresholds.

$$\text{True positive rate} = \frac{TP}{TP + FN}, \text{False positive rate} = \frac{FP}{TN + FP}, \tag{S4}$$

The MCC (defined in Equation S5) was computed due to its suitability for highly imbalanced classification tasks—such as in our case, where non-IBS regions outnumber IBS regions on protein surfaces. An MCC score ranges from –1 to +1, where +1 indicates perfect classification, 0 suggests random guessing, and –1 indicates perfectly incorrect classification. MCC has also been employed in previous benchmark comparisons of IBS predictors.[13, 14] We evaluated model performance at both the surface-level and residue-level by computing per-protein ROC AUC and MCC scores. For the benchmark comparison, since MaSIF-PMP outputs continuous prediction scores while MCC is calculated from discrete counts of TP, TN, FP, and FN, the best MCC value for each protein was determined across all tested thresholds.

$$\text{MCC} = \frac{TP \cdot TN - FP \cdot FN}{\sqrt{(TP + FP)(TP + FN)(TN + FP)(TN + FN)}} \tag{S5}$$

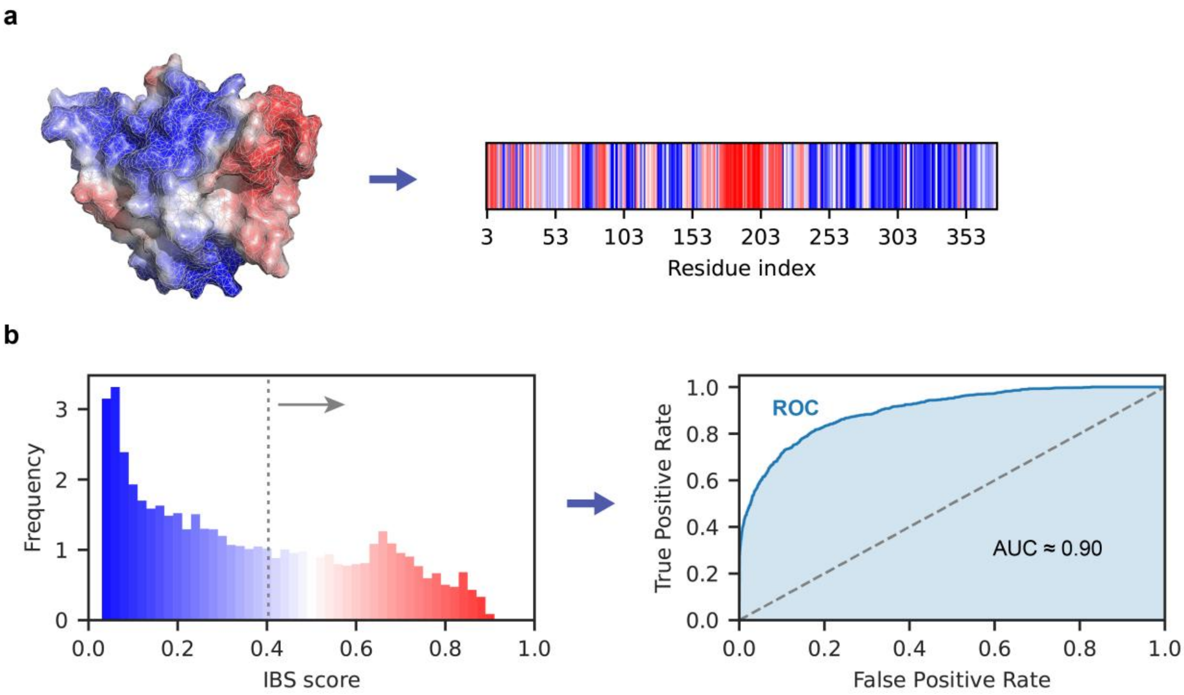


**Supplementary Figure 5.** Benchmark comparison with IBS predictors. **(a)** Schematic illustration of the procedure used to map surface-level predictions to per-residue scores. For each residue, the highest MaSIF-PMP IBS score among its associated surface points was assigned. Snapshots shown correspond to the prediction results for (S)-mandelate dehydrogenase (PDB ID: 6BFG). **(b)** Example ROC curve generated using surface-level predictions for (S)-mandelate dehydrogenase (PDB ID: 6BFG) from the benchmark set. The distribution of surface-level IBS scores for the protein is shown, with the gray dashed line indicating the sliding threshold (0–1). Points with scores equal to or above the threshold are classified as IBSs, while those below are classified as non-IBSs. Corresponding true and false positive rates were computed across thresholds to construct the ROC curve, yielding an area under the curve (AUC) of 0.90.

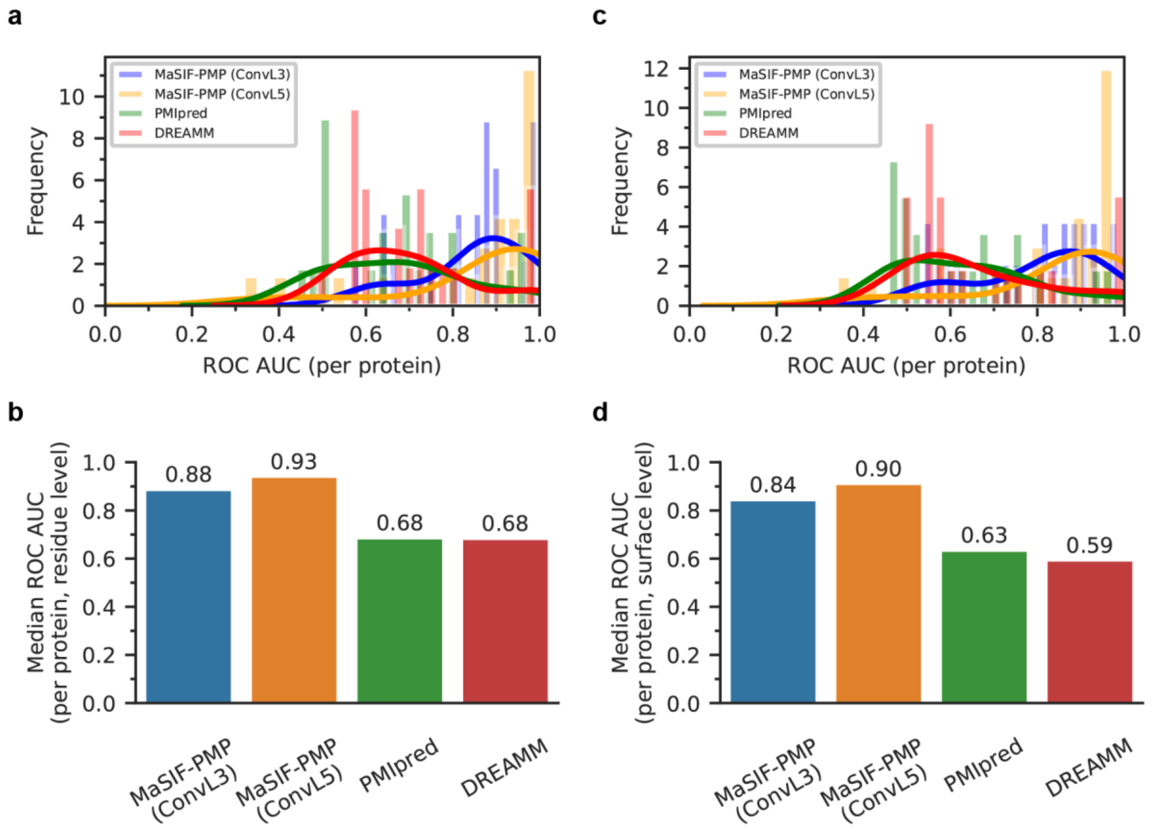


**Supplementary Figure 6.** Benchmark comparison between different IBS predictors in terms of ROC AUC scores for 21 single-chain PMPs. For MaSIF-PMP, “ConvL3” is the network with three convolutional layers while “ConvL5” is the model with five layers. **(a)** Distribution of per-protein, residue-level ROC AUC predicted by MaSIF-PMP, PMIpred[13], and DREAMM[14]. Solid lines represent Gaussian kernel density estimates fitted to the discrete score distributions. **(b)** Comparison of MaSIF-PMP with other IBS predictors on the benchmark proteins. Results are reported as the median ROC AUC per protein, evaluated on a per-residue basis. **(c–d)** Equivalent results for per-protein, surface-level ROC AUC.

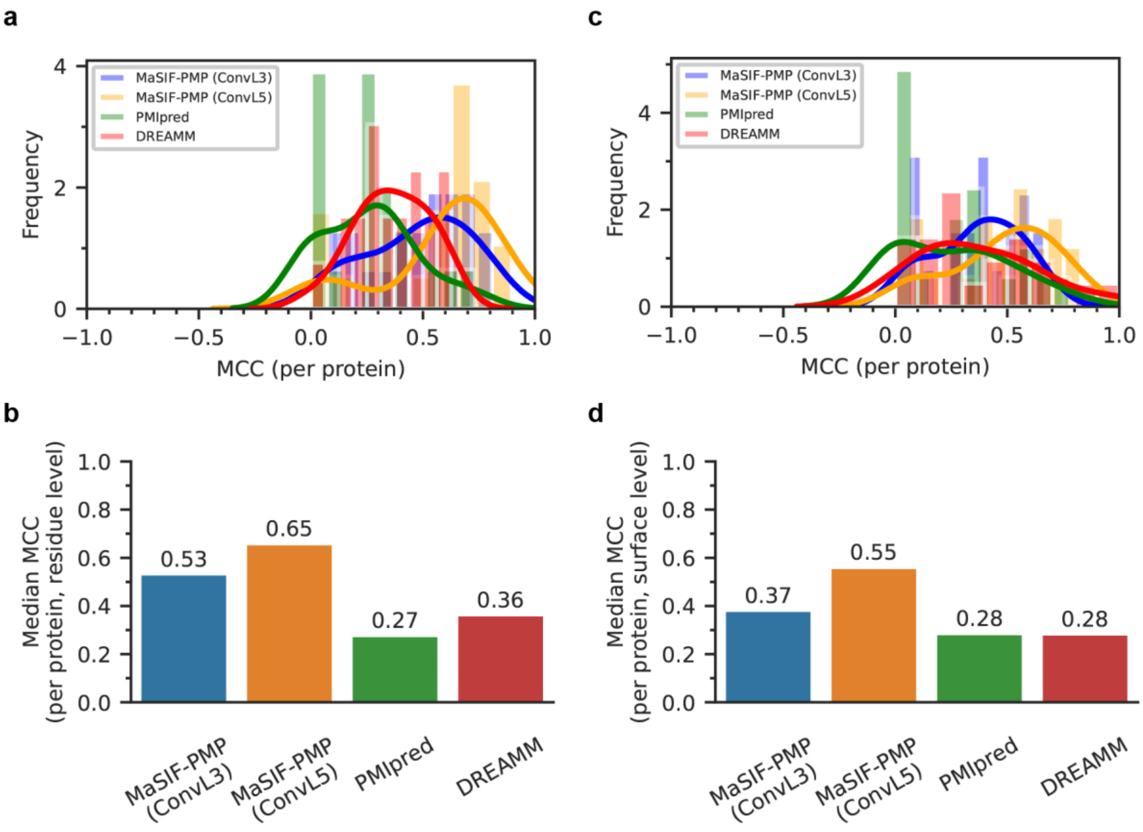


**Supplementary Figure 7.** Benchmark comparison between different IBS predictors in terms of MCC scores. The same predictor labels as in Supplementary Fig. 6 are used. **(a)** Distribution of per-protein, residue-level MCC predicted by multiple IBS predictors, with solid lines representing Gaussian kernel density estimates fitted to the discrete score distributions. For MaSIF-PMP, the best per-protein MCC was determined across all thresholds. **(b)** Comparison of MaSIF-PMP with other IBS predictors on the benchmark proteins, reported as the median per-protein MCC values evaluated on a per-residue basis. **(c–d)** Equivalent results for per-protein, surface-level MCC.

### Binary predictions of MaSIF-PMP using optimal thresholds

While MaSIF-PMP outputs continuous prediction scores representing the confidence that a surface patch belongs to an IBS, binary labels—comparable to those from other IBS predictors[13, 14]—can be obtained by applying a threshold. To determine the optimal thresholds for residue- and surface-level predictions, we calculated the median per-protein MCC across all training set proteins for thresholds ranging from 0 to 1 using the MaSIF-PMP model with three convolutional layers. The median MCC values were plotted as functions of the threshold values, and the optimal threshold values were determined as those yielding the maximum median MCC (Supplementary Fig. 8a, 8d). Using these thresholds, MCC scores were computed for the benchmark set and visualized as score distributions (Supplementary Fig. 8b, 8e) and median values (Supplementary Fig. 8c, 8f). With binary predictions based on the optimal thresholds, MaSIF-PMP achieved better or comparable prediction performance against the other IBS predictors. Specifically, kernel density estimates of MCC showed right-shifted distributions for MaSIF-PMP compared to other predictors at both residue and surface levels, with overall higher median MCC values.

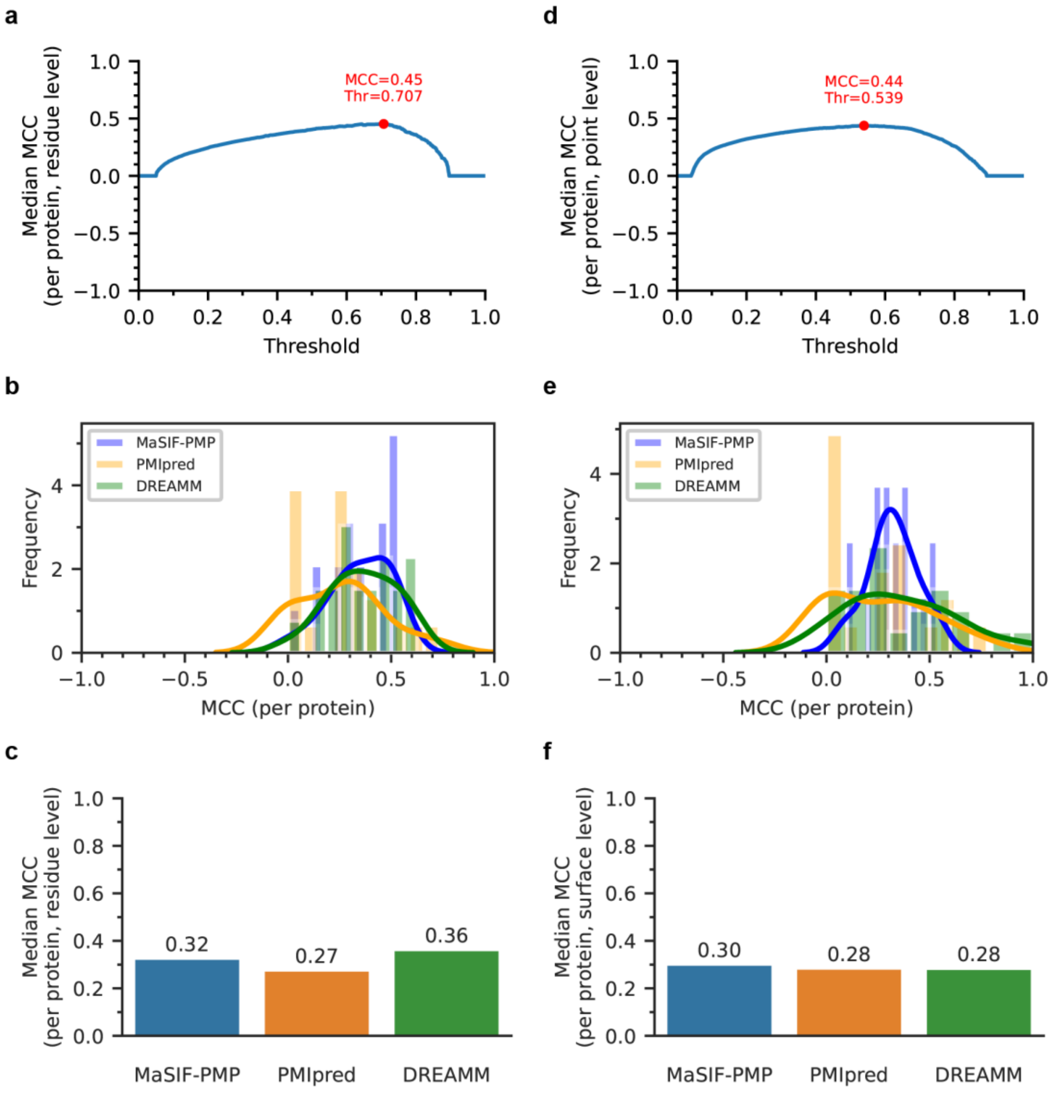


**Supplementary Figure 8**. Applying a threshold to generate binary predictions of MaSIF-PMP with three convolutional layers. **(a)** Median per-protein, residue-level MCC as a function of threshold values. Per-protein MCC values were calculated across all residues of proteins in the training set for thresholds ranging from 0 to 1, with the maximum median MCC observed at a threshold of 0.707. **(b)** Distributions of per-protein, residue-level MCC predicted by MaSIF-PMP, PMIpred, and DREAMM, using a threshold of 0.707 for MaSIF-PMP predictions. Solid lines represent Gaussian kernel density estimates fitted to the discrete score distributions. **(c)** Comparison of MaSIF-PMP using the same threshold with other IBS predictors on the benchmark proteins. Results are reported as the median MCC per protein, evaluated on a per-residue basis to ensure comparability across predictors. **(d–f)** Equivalent results for per-protein, surface-level MCC. Surface-level binary predictions of MaSIF-PMP were generated using a threshold of 0.539.

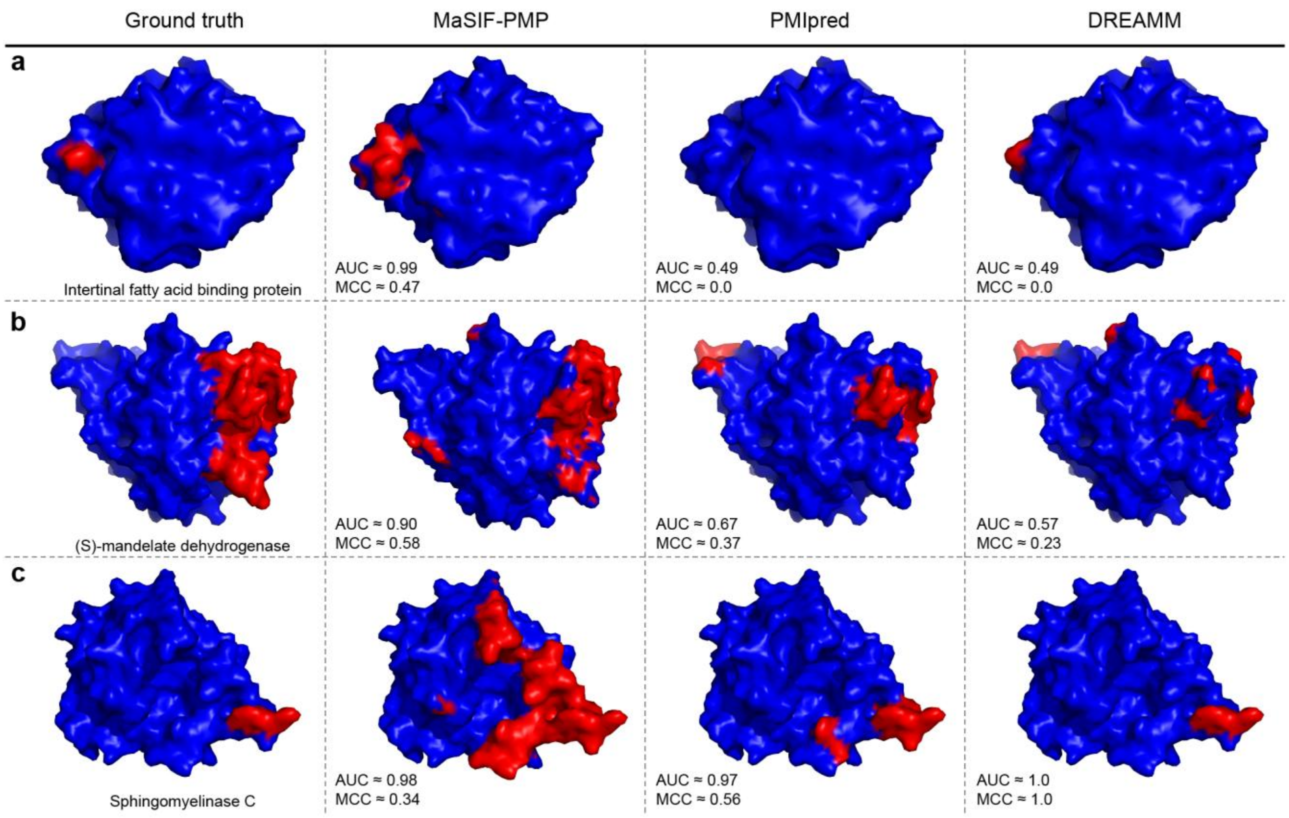


**Supplementary Figure 9**. Visualization of ground-truth and predicted IBSs for three PMPs selected from the benchmark test set. Each column shows PMP molecular surfaces with colors indicating binary labels (blue for non-binding surface points or red for the IBS) for the ground-truth, MaSIF-PMP, PMIpred, and DREAMM columns. A threshold IBS score of 0.707 (Supplementary Fig. 8) was applied to convert the continuous interface prediction scores for MaSIF-PMP into binary predictions. ROC AUC and MCC values were computed based on surface-level predictions for all three models. **(a)** Intestinal fatty acid binding protein (PDB ID: 3AKM). **(b)** (S)-mandelate dehydrogenase (PDB ID: 6BFG). **(c)** Sphingomyelinase C (PDB ID: 2DDR).

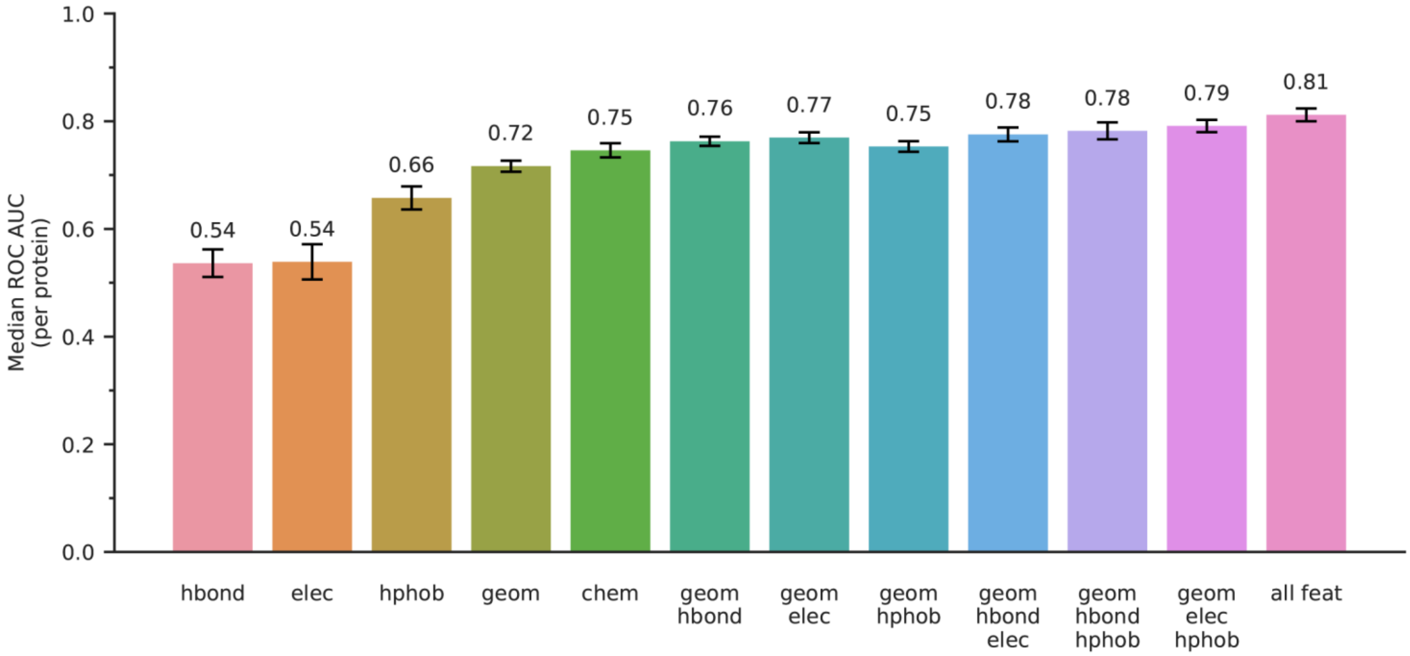


**Supplementary Figure 10**. Ablation studies using 5-fold cross-validation with MaSIF-PMP trained on different subsets of surface features: location of free electrons/proton donors (hbond), Poisson–Boltzmann electrostatic potential (elec), hydropathy index (hphob), geometric features only (geom), chemical features only (chem), and all features combined (all feat). Combinations of geometric and chemical features are denoted using the corresponding abbreviations (*e.g.*, “geom hbond” indicates training on geometric features and hydrogen bond potential).

### Transfer learning of PPI-trained model to IBS predictions

We trained the MaSIF-site using the architecture, training set, and parameters used in the previous work.[2] For transfer learning strategy 1, we froze the parameters of the three convolutional layers and replaced the final multilayer perceptron (MLP) blocks with a deeper one: a fully connected network (FCN) of FC128, FC64, FC4, FC2. For transfer learning strategy 2, we froze the parameters of the three convolutional layers then added three new convolutional layers before the final MLP blocks. Both models were then trained for 50 epochs with all 5 surface features using the same soft grid parameters and data sets introduced in *Method* section for MaSIF-PMP. The schematics of model architectures of each transfer learning model are shown in Supplementary Fig. 11.

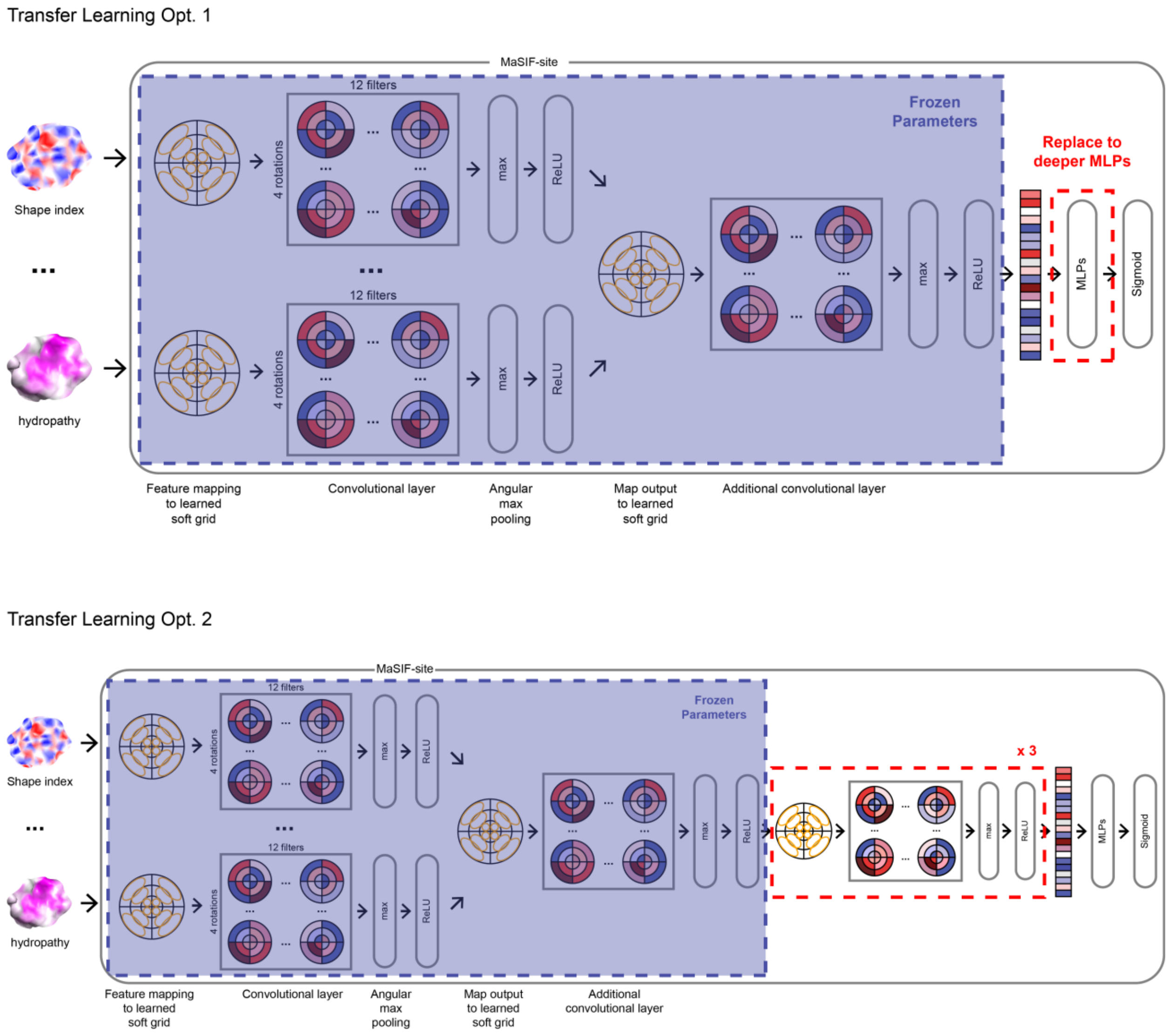


**Supplementary Figure 11**. Network architectures of MaSIF-PMP with two different transfer learning strategies that derive models from the MaSIF-site model, which is trained for PPIs. The blue shaded regions represent frozen layers where parameters trained for PPIs are no longer updated.

### Data augmentation using MD simulations of PMPs in aqueous solution

High-throughput simulations of PMPs in aqueous solution were performed using high-throughput MD (HTMD).[16] From the training set, 503 proteins were selected for the simulations based on the absence of unmodeled or missing residues within the central regions of their sequences, as such gaps can result in trajectories with multiple disconnected segments freely diffusing in solution. Protein structures were protonated at pH 7.0, with standard charged termini (*i.e.*, N-terminus: $–NH_3^+$; C-terminus: $–COO^-$). Systems were solvated with TIP3P water and neutralized using $Na^+$ and $Cl^−$ ions at 0 M concentration. All systems were parameterized using the CHARMM36m force field.[17, 18] Each system underwent energy minimization using the conjugate gradient algorithm for 500 steps, followed by a 1 ns equilibration. Production simulations were then carried out for 10 ns under constant NPT conditions. The temperature was maintained at 298.15 K using a Langevin thermostat, and pressure was maintained at 1 bar using an isotropic Monte Carlo barostat. Simulation snapshots were saved every 50 ps during production. After completion, all trajectories were preprocessed using GROMACS 2021[19] to center and apply rotational and translational fitting of the protein.

Dynamic conformations of proteins sampled from simulations were clustered using CLoNe[20], an automated clustering algorithm based on principal component analysis (PCA) of the Cartesian coordinates of $C_\alpha$ atoms. The method first performs a nearest-neighbor step to estimate local densities for each data point, followed by identification of putative cluster centers as local density maxima. Clusters are merged, if necessary, using the Bhattacharyya coefficient[21], and outliers are removed via a Bayes classifier. CLoNe requires only a single user-defined parameter, $p_{dc}$, which determines the number of clusters. The value of $p_{dc}$ can be increased to reduce the number of clusters or decreased to increase it; integer values between 1 and 10 are generally sufficient, with many values yielding identical results. In this study, the default value of 4 was used for $p_{dc}$. After clustering protein conformations sampled from HTMD simulations using CLoNe, a total of 200 conformations were grouped into representative clusters of varying sizes. To ensure training on only “meaningful” conformational states, cluster centers were filtered based on cluster size, retaining only those containing at least 5% of the total simulation frames (≥10 frames) across the entire trajectory.

### Ensemble learning with MaSIF-PMP and MD simulation data

We trained a version of the MaSIF-PMP network using an MD-augmented dataset, in which each protein was represented by multiple conformations sampled from HTMD simulations and clustered using the CLoNe algorithm. In this initial implementation, each conformation derived from a single PDB structure was treated as an independent input during training. No weighted averaging or feature aggregation across conformational ensembles was applied. As a result, the size of the training dataset increased from 1,059 to 1,896 protein structures. Training followed the same protocol as for the baseline MaSIF-PMP model. Training was performed for 100 epochs on an NVIDIA L40 GPU. The model was saved whenever the validation ROC AUC improved, with the final model checkpoint corresponding to epoch 61. No MD-based data augmentation was applied to proteins in the test set.

Despite the increased training data, the MD-augmented model exhibited similar predictive performance to the original model trained exclusively on crystal structures. On the test set, the MD-augmented network achieved a mean per-protein ROC AUC of 0.77 and a median of 0.78, compared to 0.76 (mean) and 0.78 (median) for the baseline model.

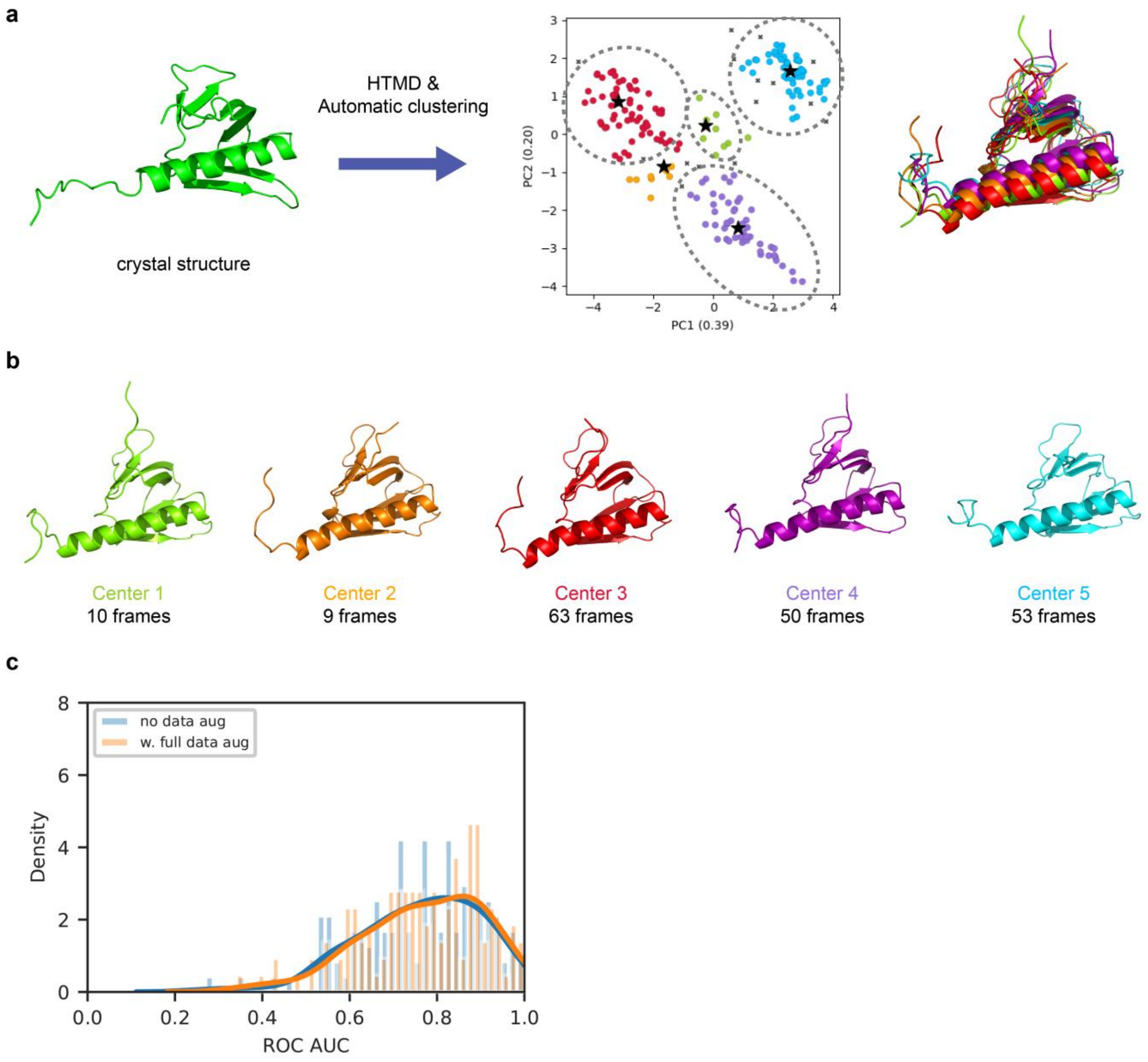


**Supplementary Figure 12.** Data augmentation using MD simulations. **(a)** Schematic overview of the workflow and concept for using MD simulations to capture protein conformational dynamics. All snapshots and plots are based on G-protein coupled receptor kinase 2 (PDB ID: 1BAK). Gray dashed circles highlight representative cluster centers selected from the trajectory based on frame counts. The ensemble snapshot on the right shows the superposition of cluster center structures. **(b)** Representative cluster centers sampled from the HTMD trajectory using the CLoNe clustering algorithm. **(c)** Distribution of per-protein surface-level ROC AUC scores across the test set, predicted by MaSIF-PMP models trained with and without MD-based data augmentation. Kernel density estimates were generated using Gaussian kernels.

### Case studies using HMMM simulations

α-tocopherol transfer protein (α-TTP)

HMMM simulations of α-tocopherol transfer protein (α-TTP) were conducted using the crystal structure from PDB ID: 3W67, chain A. Simulations were performed with α-TTP in complex with α-tocopherol (α-Tol) because α-TTP binds to membranes in its ligand-bound state and undergoes a ligand-exchange mechanism between phosphatidylinositol phosphates (PIPs) and α-Tol.[22] Although α-Tol is buried within the hydrophobic core of the protein and has minimal effect on the molecular surface, it was included to accurately model the biologically relevant conformation.

Membrane binding in α-TTP occurs primarily via direct interactions between the negatively charged $PIP_2$ headgroups and basic residues located near the opening of the ligand-binding cavity.[22] The HMMM membrane system was constructed to replicate the lipid composition used in prior MD study[22] of α-TTP, consisting of 60 DOPC, 40 DOPE, and 2 $PI(4,5)P_2$ molecules per leaflet, with DCLE molecules modeling the hydrophobic core (Supplementary Table 3). For replica simulations (Supplementary Fig. 13a), we tested three systems with different orientations of α-TTP with respect to the HMMM membrane: the same orientation of structure from RCSB PDB (*ori.1*), rotated along the $x$ axis by 90° (*ori.2*), and rotated along the $y$ axis by 90° (*ori. 3*). We performed three replica simulations with different initial velocities for each orientation for a total of 9 independent simulations. Detailed system information is provided in Supplementary Table 3.

All systems were energy-minimized using the steepest descent algorithm until the maximum force between atoms reached the criterion of <1000 kJ $mol^{-1}$ $nm^{-2}$. After energy minimization, systems were equilibrated at constant NVT for 250 ps then further equilibrated for 1,625 ps at constant NPT. Positional restraints on HMMM lipid atoms were gradually reduced during equilibration. Production simulations were performed for 100 ns under NPT conditions at 300 K using a velocity-rescale thermostat (time constant = 1.0 ps) and at 1 bar using a semi-isotropic stochastic cell-rescaling barostat (time constant = 5.0 ps, compressibility = $4.5 \times 10^{-5}$ $bar^{-1}$). All MD simulations were conducted with a 2 fs timestep using the leapfrog integrator in Gromacs 2021.[19] Verlet lists were generated with a 1.2 nm neighbor list cutoff, van der Waals interactions were modeled using a Lennard-Jones potential with a 1.2 nm cutoff that was smoothly shifted to zero between 1.0 and 1.2 nm, and electrostatic interactions were calculated using the smooth particle-mesh Ewald method with a short-range cutoff of 1.2 nm.[23] Bonds involving hydrogen atoms were constrained using the LINCS algorithm.[24] Further simulation parameters and raw data are available in the published dataset (DOI: 10.5061/dryad.1rn8pk175). α-TTP exhibited stable membrane interactions only in orientation 1, with no consensus IBSs detected for other orientations (Supplementary Fig. 13); consensus IBSs are defined in the next section.

**Supplementary Table 3.** Number of components for simulated α-TTP systems.

| | | *ori.1* | *ori.2* | *ori.3* |
|---|---|---|---|---|
| α-TTP | | 1 | 1 | 1 |
| Lipid molecules | $PI(4,5)P_2$ | 4 | 4 | 4 |
| | PE | 80 | 80 | 80 |
| | PC | 120 | 120 | 120 |
| | DCLE | 1,192 | 1,196 | 1,192 |
| α-Tol | | 1 | 1 | 1 |
| Water molecules | | 23,202 | 21,528 | 22,331 |
| $Na^+$ ions | | 14 | 14 | 14 |
| $Cl^-$ ions | | 0 | 0 | 0 |
| Total atoms | | 97,130 | 92,140 | 94,517 |

Defining consensus and union IBS label based on trajectories of replica simulations

Using replica simulations with multiple protein orientations relative to the membrane, we defined consensus IBS labels based on the fraction of simulation time during which protein–membrane contacts were observed. The final 10 ns of each replica were used to define these labels because stable contacts were consistently observed in this interval. Consensus IBS regions were designated based on the residues within 0.5 nm of any lipid (except the DCLE molecules that escaped from the bilayer) atom for at least 90% of the aggregated production phases (Supplementary Fig. 13b). Union IBS labels were defined by combining consensus IBS labels for each orientation (Supplementary Fig. 13c).

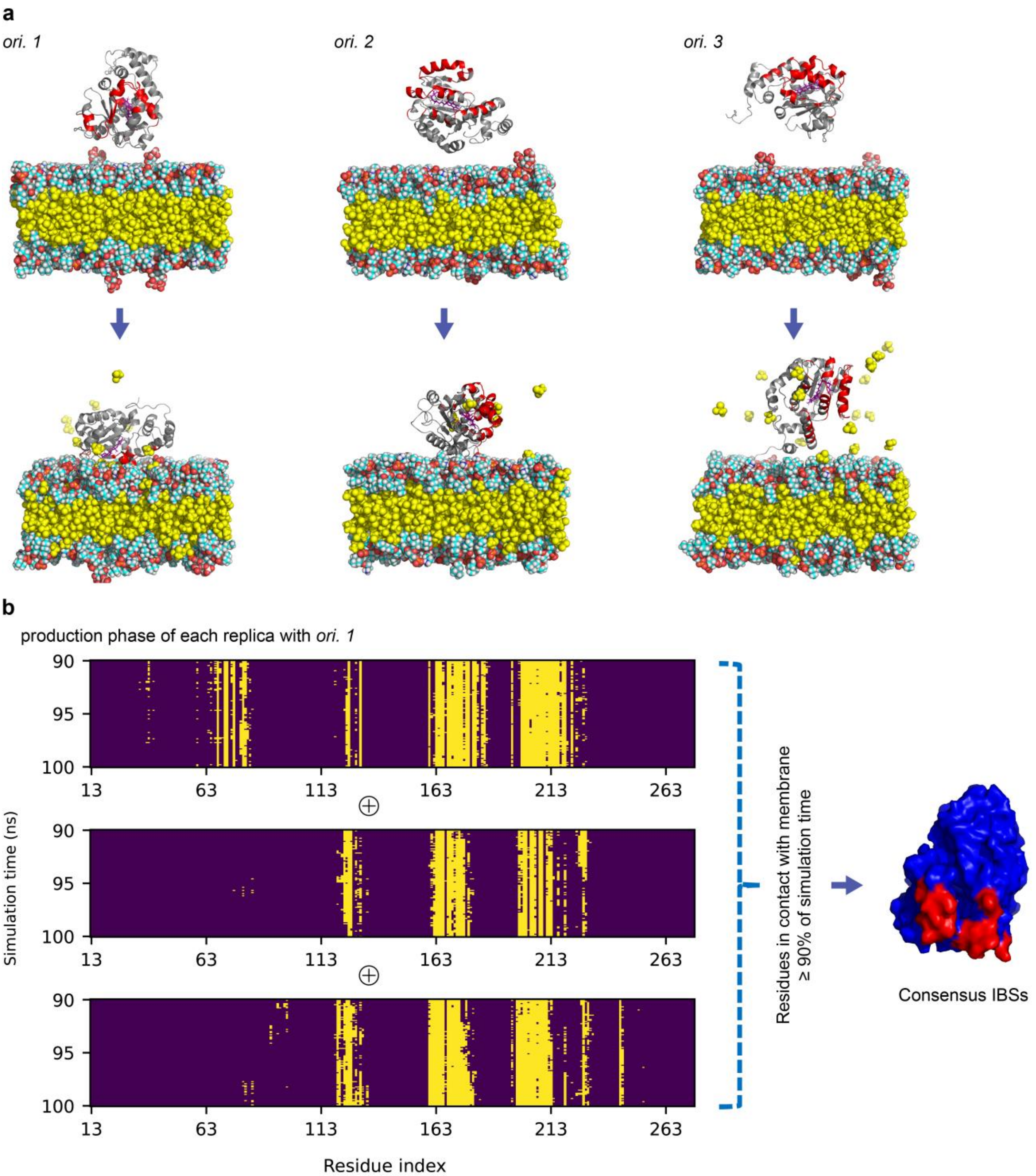


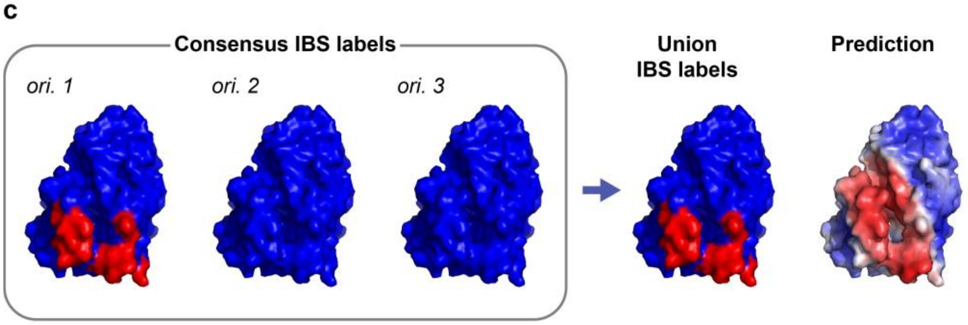


| Consensus Label | ROC AUC |
|---|---|
| *ori. 1* | 0.83 |
| *ori. 2* | N/A |
| *ori. 3* | N/A |
| union | 0.83 |

**Supplementary Figure 13.** Replica simulations of α-TTP to check the consistency and robustness of IBS labels determined based on HMMM MD. Replica simulations were performed with α-TTP in multiple orientations with respect to HMMM membrane. **(a)** Representative snapshots of α-TTP before/after 100 ns HMMM MD simulations across multiple initial orientations (*ori.*). Snapshots of replica 1 for each orientation are presented. The IBS region derived from replica 1 of orientation 1 is highlighted in red for comparison between simulations with different orientations. The HMMM membrane is shown in a van der Waals representation; carbon atoms in cyan, hydrogen atoms in white, oxygen atoms in red, nitrogen atoms in blue, phosphorus atoms in orange, α-tocopherol in purple, and organic solvent 1,1-dichloroethane (DCLE) in yellow. Water molecules and ions are not shown to aid visualization. **(b)** Defining consensus IBSs based on the fraction of simulation time in which protein–membrane contacts were observed. The final 10 ns of each replica were considered as the production phase, and we applied a 90% cutoff to designate consensus IBS regions. Only results from orientation 1 (*ori.1*) are shown in the figure. For the barcode plots, purple indicates no contact while yellow indicates contact between the protein and membrane. **(c)** Snapshots of consensus IBS labels for each orientation, their union, and predicted IBS scores for α-TTP. Tables show corresponding ROC AUC values computed with different ground-truth label types.

Oxysterol-binding protein homologue (Osh4)

HMMM simulation of oxysterol-binding protein homologue (Osh4) was performed using the crystal structure from PDB ID: 1ZHZ (chain A). Osh4 is known to undergo conformational rearrangement when interacting with trans-Golgi network (TGN) anionic membrane. A TGN membrane was prepared using the same lipid compositions as in a prior MD study of Osh4 (Supplementary Table 4).[25] The system was neutralized with NaCl counterions.

MD simulations were performed for 100 ns under NPT conditions at 303.15 K and 1 bar, using the velocity-rescale thermostat and semi-isotropic stochastic cell-rescaling barostat. The same parameters as in the simulation of α-TTP were used for Osh4 simulations. Further simulation parameters and raw data are available in the published dataset (DOI: 10.5061/dryad.1rn8pk175). For the ROC AUC computation, we also compared HMMM labels to ground-truth labels derived from previous MD studies that reported six separated membrane-binding domains (referred to as the “paper” labels in Supplementary Fig. 14).[25, 26]

**Supplementary Table 4.** Composition of simulated system for the oxysterol-binding protein homologue (Osh4). Acronyms: TGN, trans-Golgi network; ERG, ergosterol; DYPC, PC(16:1(9Z)/16:1(9Z)); DYPE, PE(16:1(9Z)/16:1(9Z)); POPA, PA(16:0/18:1(9Z)); POPI, PI(16:0/18:1(9Z)); POPS, PS(16:0/18:1(9Z)); PYPE, PE(16:0/16:1(9Z)); PYPI, PI(16:0/16:1(9Z)); YOPC, PC(16:1(9Z)/18:1(9Z)); YOPE, PE(16:1(9Z)/18:1(9Z)); DCLE, 1,1-dichloroethane. For lipid names, the first two characters indicate the headgroup type, while the numbers and "Z" notation denote the acyl chain length and position of the cis double bond, respectively.

| | | Number |
|---|---|---|
| Osh4 | | 1 |
| TGN | ERG | 36 |
| | DYPC | 58 |
| | DYPE | 12 |
| | POPA | 8 |
| | POPI | 54 |
| | POPS | 10 |
| | PYPE | 8 |
| | PYPI | 56 |
| | YOPC | 46 |
| | YOPE | 12 |
| | DCLE | 1,376 |
| Water molecules | | 36,565 |
| $Na^+$ ions | | 138 |
| $Cl^-$ ions | | 0 |
| Total atoms | | 148,753 |

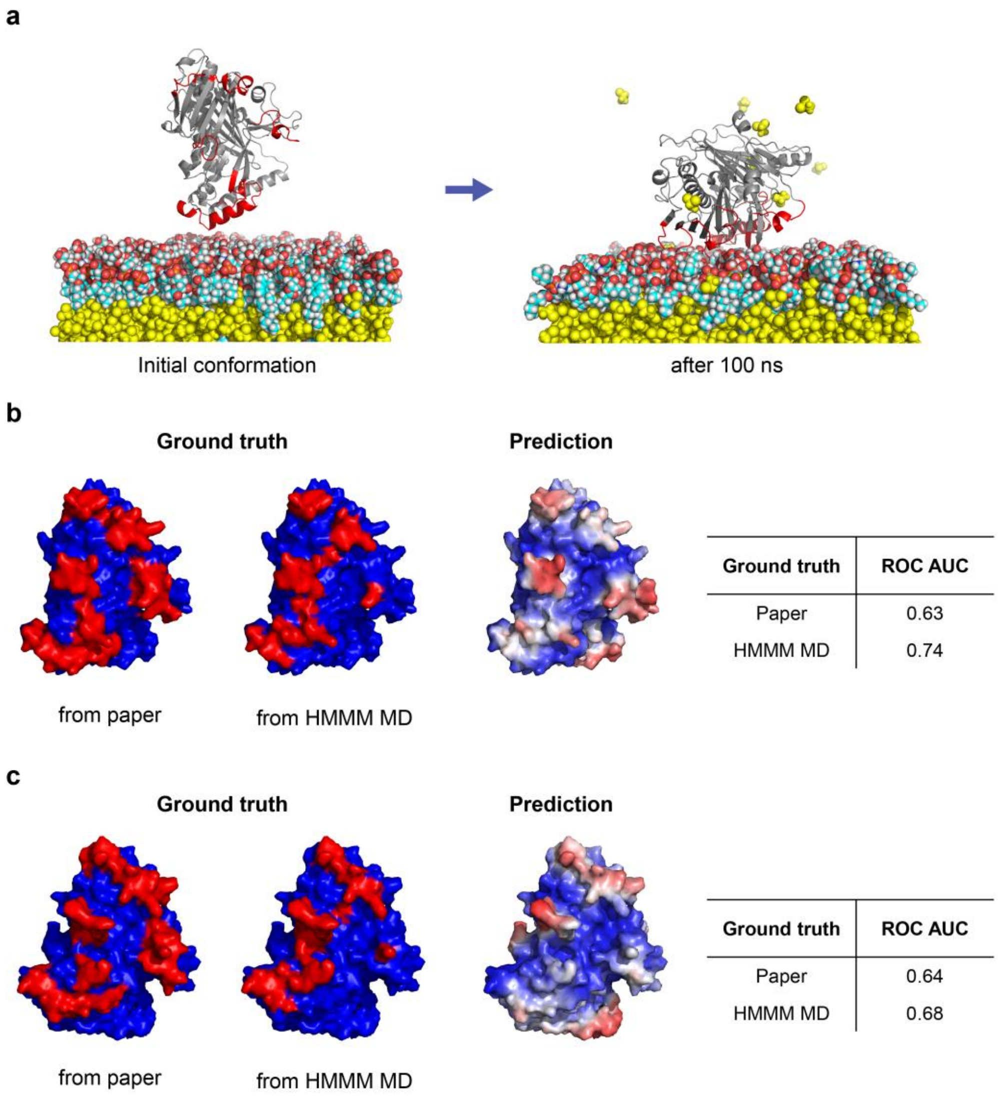


| Ground truth | ROC AUC |
|---|---|
| Paper | 0.63 |
| HMMM MD | 0.74 |

| Ground truth | ROC AUC |
|---|---|
| Paper | 0.64 |
| HMMM MD | 0.68 |

**Supplementary Figure 14.** Case study with Osh4 using HMMM MD simulations. **(a)** Snapshot of Osh4 (PDB ID: 1ZHZ) interacting with the trans-Golgi network (TGN) membrane, sampled from HMMM MD simulations. Ground-truth membrane-binding interfaces from prior studies[25, 26] are highlighted in red. Color schemes follow the same definition used in Supplementary Figure 13a. **(b)** MaSIF-PMP prediction on the crystal structure of Osh4. Reported ROC AUC scores correspond to surface-level predictions. ROC AUC values are computed using ground-truth labels from either previous studies (referred to as the "paper" ground-truth labels) or from the HMMM MD simulations. **(c)** MaSIF-PMP prediction on the membrane-bound conformation of Osh4 captured from HMMM simulations. Reported ROC AUC values are also based on surface-level predictions.

Representative HMMM membrane types: anionic and zwitterionic

To investigate membrane-binding behaviors of PMPs with poor MaSIF-PMP prediction performance, we selected two representative membrane types: anionic and zwitterionic. These types reflect distinct chemical characteristics commonly observed in biological systems, such as of the trans-Golgi network (TGN) and endoplasmic reticulum (ER), two well-studied membranes known for their differing lipid compositions and surface chemistries.[27] Although both biological membranes consist of a mix of zwitterionic and anionic lipids, we simplified the model systems by including only zwitterionic phosphatidylethanolamine (PE) and anionic phosphatidylserine (PS). This simplification was guided by the widespread biological relevance of PE and PS, as well as their comparable lengths, which help minimize geometric variations in the membrane surface. Accordingly, the HMMM membranes were prepared using the following lipid ratios: PE:PS = 100:0 for the zwitterionic membrane, and PE:PS = 50:50 for the anionic membrane. For the replica simulations, three initial protein orientations with respect to the HMMM membrane were tested, identical to those used for the α-TTP systems. A single replica simulation was performed for each orientation.

The structure of phospholipase A2 was obtained from PDB ID: 1OZY. Additional disulfide bonds were modeled between the following cysteine pairs: C11-C77, C29-C45, C44-C105, C51-C98, C61-C91, and C84-C96. Detailed system information is provided in Supplementary Table 5. Further simulation parameters and raw data are available in the published dataset (DOI: 10.5061/dryad.1rn8pk175).

The structure of glycosyl hydrolase was obtained from PDB ID: 4LPL. As the net charge of the glycosyl hydrolase system with the zwitterionic HMMM membrane was zero, the system was neutralized using 0.02 M NaCl. Detailed system information is provided in Supplementary Table 5. Further simulation parameters and raw data are available in the published dataset (DOI: 10.5061/dryad.1rn8pk175).

We defined the consensus IBS labels of the two proteins with either HMMM membrane types in the same manner used for α-TTP system in the previous section. We ran replica simulations for each initial orientation, used the final 10 ns of each replica as the production phase, and applied a time-fraction cutoff of 95% to define the consensus IBS labels for each orientation. The union of the HMMM-derived IBS labels for each replica was compared with the MaSIF-PMP predictions to compute the ROC AUC values reported in the main text.

**Supplementary Table 5.** Number of components for each simulated system. The final composition of each system is summarized below. PE denotes phosphatidylethanolamine, PS denotes phosphatidylserine, and DCLE refers to the organic solvent 1,1-dichloroethane.

| | | | PE | PS | DCLE | Water molecules | $Na^+$ ions | $Cl^-$ ions | Total atoms |
|---|---|---|---|---|---|---|---|---|---|
| Phospholipase A2 | Anionic membrane | *ori.1* | 100 | 100 | 1,008 | 19,157 | 99 | 0 | 79,870 |
| | | *ori.2* | 100 | 100 | 1,008 | 18,664 | 99 | 0 | 78,391 |
| | | *ori.3* | 100 | 100 | 986 | 16,934 | 99 | 0 | 73,025 |
| | Zwitterionic membrane | *ori.1* | 200 | 0 | 1,004 | 19,079 | 0 | 1 | 79,306 |
| | | *ori.2* | 200 | 0 | 1,000 | 18,562 | 0 | 1 | 77,723 |
| | | *ori.3* | 200 | 0 | 1,004 | 16,837 | 0 | 1 | 72,580 |
| Glycosyl hydrolase | Anionic membrane | *ori.1* | 100 | 100 | 998 | 18,026 | 100 | 0 | 76,842 |
| | | *ori.2* | 100 | 100 | 1,008 | 17,985 | 100 | 0 | 76,799 |
| | | *ori.3* | 100 | 100 | 998 | 17,755 | 100 | 0 | 76,029 |
| | Zwitterionic membrane | *ori.1* | 200 | 0 | 1,004 | 17,914 | 6 | 6 | 76,266 |
| | | *ori.2* | 200 | 0 | 1,004 | 17,924 | 6 | 6 | 76,296 |
| | | *ori.3* | 200 | 0 | 991 | 17,674 | 6 | 6 | 75,442 |

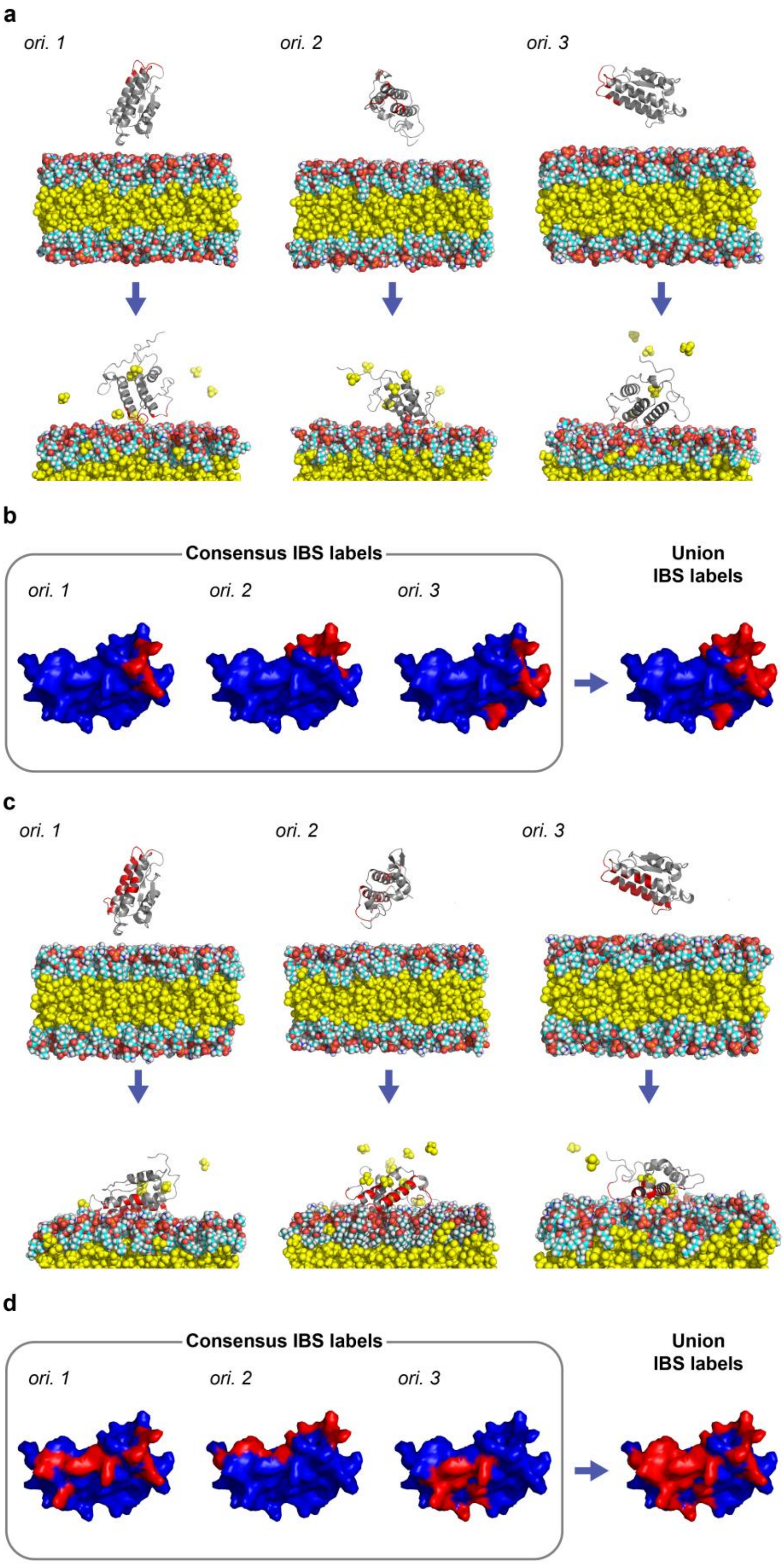
a
ori. 1
ori. 2
ori. 3
b
Consensus IBS labels
ori. 1
ori. 2
ori. 3
Union
IBS labels
c
ori. 1
ori. 2
ori. 3
d
Consensus IBS labels
ori. 1
ori. 2
ori. 3
Union
IBS labels

**Supplementary Figure 15.** Replica simulations of the phospholipase A2 (PDB ID: 1OZY) system to check the consistency and robustness in IBS labels determined based on HMMM MD. Replica simulations were performed with the protein initialized in multiple orientations with respect to HMMM membrane. **(a)** Representative snapshots of phospholipase A2 after 100 ns HMMM MD simulation with the anionic membrane across multiple initial orientations (*ori.*). IBS regions derived from replica 1 (*ori.1*) is highlighted in red for comparison across orientations. Color schemes follow the same definition used in Supplementary Figure 13a. **(b)** Snapshots of consensus IBS labels for each orientation and their union derived from HMMM simulations with the anionic membrane. Consensus IBSs were defined based on the fraction of simulation time in which protein–membrane contacts were observed, using the final 10 ns of each replica as the production phase and a 95% cutoff for designation. **(c)** Representative snapshots of phospholipase A2 after 100 ns of HMMM MD with the zwitterionic membrane, and **(d)** the corresponding consensus IBS labels for each orientation and their union, presented in the same manner as panels (a) and (b).

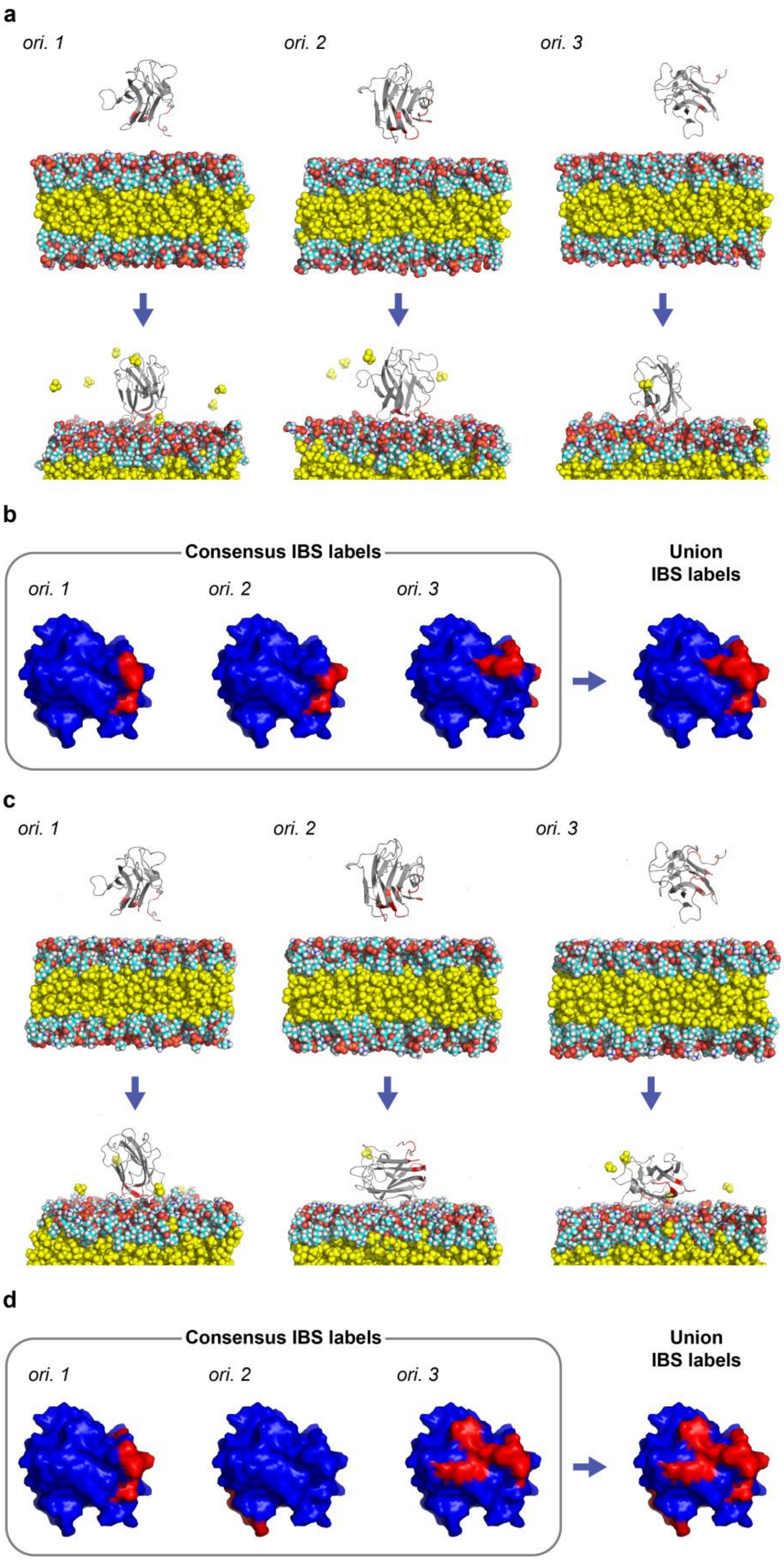
a
ori. 1
ori. 2
ori. 3
b
Consensus IBS labels
ori. 1
ori. 2
ori. 3
Union
IBS labels
c
ori. 1
ori. 2
ori. 3
d
Consensus IBS labels
ori. 1
ori. 2
ori. 3
Union
IBS labels

**Supplementary Figure 16.** Replica simulations of the glycosyl hydrolase (PDB ID: 4LPL) system to check the consistency and robustness in IBS labels determined based on HMMM MD. Replica simulations were performed with the protein initialized in multiple orientations with respect to HMMM membrane. **(a)** Representative snapshots and **(b)** corresponding consensus IBS labels from 100 ns HMMM MD simulations with the anionic membrane, presented in the same manner as Supplementary Fig. 15a-b. **(c)** Equivalent snapshots and **(d)** consensus IBS labels from simulations with the zwitterionic membrane.
